# Divergence in a stress regulatory network underlies differential growth control

**DOI:** 10.1101/2020.11.18.349449

**Authors:** Ying Sun, Dong-Ha Oh, Lina Duan, Prashanth Ramachandran, Andrea Ramirez, Anna Bartlett, Maheshi Dassanayake, José R. Dinneny

## Abstract

The phytohormone abscisic acid (ABA) is a central regulator of acclimation during environmental stress. While some plants exhibit tremendous stress resilience, it has been unclear whether differences in ABA response underlie such adaptations. Here we establish a cross-species gene regulatory network (GRN) for ABA to identify broadly conserved, core components of the ABA signaling network and peripheral pathways exhibiting species-specific connectivity. Genes that are broadly conserved in the network share promoter architecture and patterns of gene expression. Networks associated with growth hormones exhibited highly divergent wiring of their ABA network leading to changes in the physiological outcome of signaling. Together our study provides a model for understanding how GRN subcircuits deploy different growth regulatory states across ecologically diverse species.

**One Sentence Summary:** Comparative studies reveal core and peripheral stress-mediated gene networks driving divergent growth control in plants.

## Main Text

The most widely characterized plants are typically sensitive to stress, thus limiting our knowledge of the most effective response mechanisms that exist, and our ability to design innovative engineering strategies to maintain agricultural yields in the face of climate change (*1*-*3*). Extremophytes are plant species that have evolved mechanisms to cope with severe abiotic stresses such as salinity, drought, and cold (*4, 5*). Within the Brassicaceae, two emerging model extremophyte species exhibit greater tolerance to soil salinity than the well-characterized plant *Arabidopsis thaliana* (*A. thaliana*) (*6, 7*). *Eutrema salsugineum* (*E. salsugineum*) is found from the saline soils in coastal China to the subarctic regions in Canada, and survives stress through effective energy use, regulation of excess electron flow, and presence of protective barriers in the root (*8*-*10*), while *Schreinkiella parvula* (*S. parvula*) grows in the Irano-Turanian region, is tolerant to high levels of Na^+^, K^+^, Li^+^, Mg^2+^, and Boron (*11, 12*), but the adaptive strategies used are less-well characterized. *S. parvula* is believed to have evolved salinity tolerance independently of *E. salsugineum* as the closely related species *Sisymbrium irio* (*S. irio*) together with the entire clade of *Brassica* are also stress sensitive (Fig. 1A) (*13*).

**Fig. 1.**
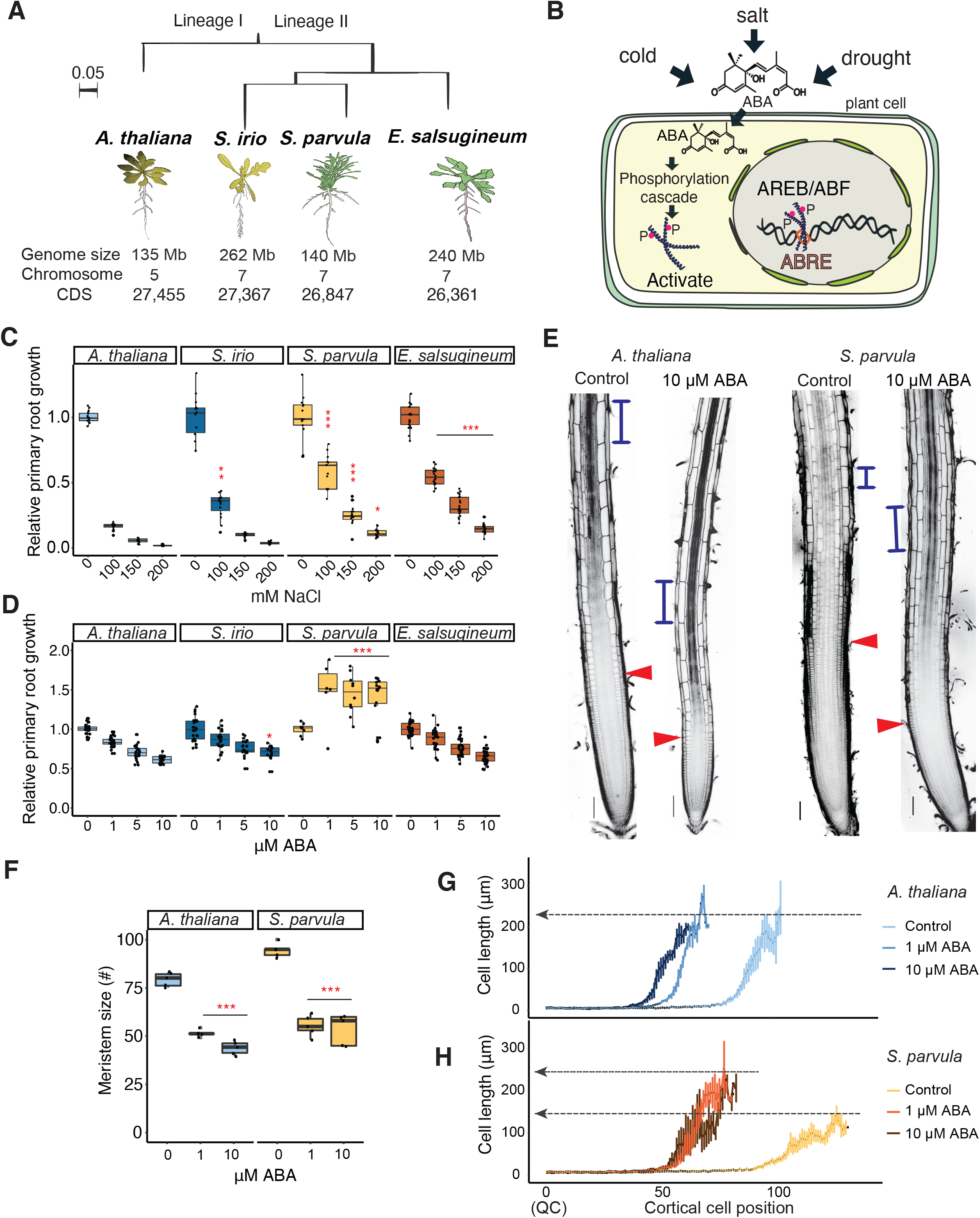
Comparative analysis of stress response in the Brassicaceae family. **(A)** A phylogenetic tree generated with lineage I, stress sensitive *A. thaliana*, lineage II stress sensitive *S. irio* and extremophytes *S. parvula* and *E. salsugineum*. Scale = amino acid per site. **(B)** Environmental stress triggers a signaling cascade and subsequent phosphorylation of bZIP TFs AREB/ABFs, which activates gene expression by binding to ABA-RESPONSIVE ELEMENTs (ABREs) (**C**,**D**) Quantification of *A. thaliana* and *S. parvula* primary roots after transfer to NaCl **(C)**, or ABA **(D)** media for 2 days. Asterisks mark significant differences from *A. thaliana* based on two-way ANOVA, \**P* < 0.05, \*\**P* < 0.01, \*\*\**P* < 0.001, n > 6. **(E)** Confocal images of primary roots grown on 10 µM ABA or standard media. Blue bars label the stabilized cortex cell length. Red arrows point to the end of the meristem, based on the root cap landmark. Scale bars = 100 µm. **(F)** Quantifications of root meristem size for *A. thaliana* and *S. parv*ula under control and 1 or 10 µM ABA treatment. Pairwise T-test, _***_*P* < 0.001, n = 5. (**G**,**H)** Quantification of cortex cell length (n = 5). The x-axis indicates cell number starting from the quiescent center (QC).

We chose to focus our comparative analysis on the response to abscisic acid (ABA) as a proxy for environmental stress as the transcriptional response to this hormone is not dependent on the organism experiencing physiological stress, which may occur under highly different environmental conditions between species (*14*). ABA biosynthesis is induced under salt stress and typically inhibits germination, root growth, and gas exchange as a means of limiting growth (*15*-*17*). ABA perception triggers a signaling cascade that leads to the phosphorylation of the bZIP transcription factor (TF), ABA-RESPONSIVE ELEMENT BINDING FACTORS (AREB/ABFs) (*18, 19*). AREB/ABFs relocalize to the nucleus and activate gene expression by binding to cis-regulatory elements (CREs) known as *ABA-RESPONSIVE ELEMENT*s (*ABREs*) (Fig. 1B).

Our growth assay confirmed the substantial tolerance of *S. parvula* and *E. salsugineum* to NaCl compared with *A. thaliana* and *S. irio* (Fig. 1C). In contrast, application of exogenous ABA had strong inhibitory effects in *A. thaliana, E. salsugineum* and *S. irio*, while *S. parvula* exhibited a striking growth promotion at all concentrations tested (Fig. 1D). We characterized the developmental basis for the observed growth enhancement (Fig. 1E) and found that while the meristem size for *A. thaliana* and *S. parvula* is reduced upon ABA treatment (Fig. 1F), final cell length for *S. parvula* increased by 50% (from ∼100 µm to ∼200 µm) while no change occurred in *A. thaliana* (Fig. 1G-H). These data reveal that all species tested respond to ABA by differential growth, but that the direction of this effect varied dramatically depending on the species and developmental context.

To identify conserved and species-specific ABA responsive pathways, we performed RNA-Seq on the roots and shoots of all 4 species treated with either mock or 10 µM ABA for 3 hours and 24 hours (data S1-S3). We characterized the number of differentially expressed genes (DEGs) and observed that *S. parvula* uniquely responded with a more sustained transcriptional response in roots and shoots, which contrasted with *A. thaliana, S. irio*, and *E. salsugineum* where the greatest number of DEGs was observed after 3 hours ABA treatment in root tissue (Fig. 2A). Similar *S. parvula*-specific trends were also observed in the prevalence of expression patterns identified after clustering (Fig. 2B).

**Fig. 2.**
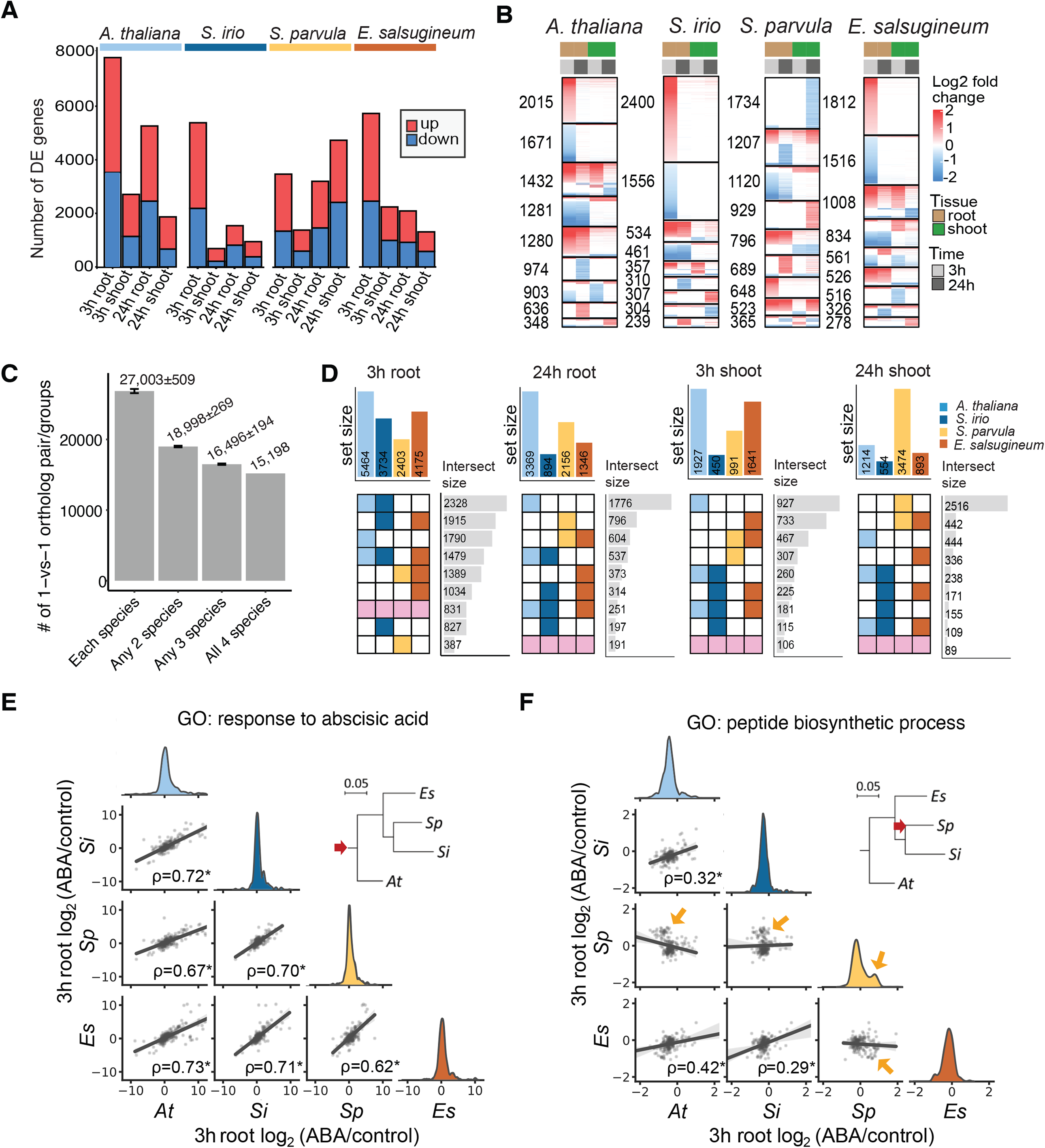
RNA-Seq identifies conserved and species-specific ABA responsive pathways. **(A)** Number of differentially expressed genes (DEGs) based on *P* <0.05. **(B)** Heatmap of Log_2_ fold changes (LFC) for DEG clusters. **(C)** Number of 1:1 orthologous groups (OGs) identified for any two, three, or all four species. Error bars indicate standard deviations. **(D)** Top 9 categories of DEGs from 1:1 OGs in descending order. The conserved responses (pink) were found to be less prevalent while responses specific to stress-sensitive plants (light and dark blue) and extremophytes (yellow and red) were most prevalent. **(E**,**F)** Phylogenetically-informed profiling (Pip) identified functional categories with lineage-specific modifications (fig. S2). Spearman’s correlations (ρ, * *Padj*<10^−7^) and linear regressions (with gray-shaded 95% confidence intervals) were shown for LFC of ortholog pairs annotated with the GO term. Red arrows mark modified lineages. “ABA-response” was conserved in all species while “peptide biosynthetic process” showed *S. parvula*-specific deviation from conserved gene expression (gold arrows). *S. parvula*-specific modifications represented the most frequent lineage-specific pattern (fig. S3a). “Peptide biosynthetic process” and related GO terms in root samples showed the most dramatic variance across species among tested GO terms (fig. S3b, table S1, and data S4).

To compare expression patterns of related genes across species, CLfinder was used to define 15,198 unambiguous 1-to-1 orthologous gene groups based on reciprocal homology (Fig. 2C, fig. S1, and data S1). Gene groups were then clustered based on their pattern of differential expression to ABA, which revealed that for each tissue and time point where expression was measured, species-specific and stress-sensitive/extremophyte-specific patterns were most prevalent while genes that respond across all species were less common (colored pink, Fig. 2D).

Leveraging the evolutionary relationships of our studied species, we developed a phylogenetically informed profiling (PiP) approach that utilizes a correlation metric to establish whether pairs of species exhibit a similar regulatory program for genes associated with distinct functional groups defined by GO terms (fig. S2 and data S4). PiP revealed that the genes annotated with the GO term “response to abscisic acid” are significantly correlated in all species pairs. These data suggest broad conservation in regulation (activated or repressed) for canonical targets of ABA signaling and that the regulatory network is derived from ancestral traits (Fig. 2E). In addition to conserved ABA response pathways, our novel analysis method revealed that orthologous gene pairs associated with protein translation (“peptide biosynthetic process”), and other related GO terms, exhibited no correlation or showed anticorrelation, between *S. parvula* and other species (Fig. 2F, fig. S2, table S1). Protein translation is finely orchestrated with the metabolic activity of cells and corresponds to the “growth” cellular state (*20*). This pattern suggests an *S. parvula*-specific change in ABA-dependent regulation that diverged from the common ancestor (Fig. 2F, fig. S3). Therefore, *S. parvula* may have evolved to interpret ABA signaling as a proxy for environmental conditions to promote growth, rather than suppress growth, contrasting to other species examined.

To determine which classes of TFs are likely involved in generating the ABA-dependent transcriptional responses observed in each species, we used Analysis of Motif Enrichment (AME), and searched for over-represented TF binding motifs within the region located 1 kb upstream of the coding sequence (*21*). We identified motifs for AREB/ABF 1/2/3/4 and 31 additional TFs, all of which included a G-box like sequence similar to the previously defined cis-regulatory element, ABRE. This motif was not observed for ABA-repressed DEGs which is in line with previous studies reporting ABREs as recruiters of transcriptional activation (*22*) (fig. S4, data S5). Our promoter analysis suggests that the molecular mechanism underlying ABA-induced gene expression is functionally conserved across species.

Based on this evidence for a shared mechanism of ABA-dependent gene expression, we sought to establish comparative global maps of AREB/ABF binding landscapes since AREB/ABFs are expressed in vegetative tissues (fig. S5) and play an indispensable and inducing role in ABA-mediated gene expression in *A. thaliana* (18, *23*-*27*). The recent development of DNA affinity purification (DAP-Seq) provided a method amendable for the study of non-model organisms (*28*). DAP-Seq facilitates the direct measurement of TF to genomic DNA (gDNA) binding without the need to develop TF-specific antibodies or generate transgenic lines with epitope-tagged TFs. Despite differences in target gene expression, the AREB/ABF gene family has not expanded or contracted, resulting in 4 homologs within each species (fig. S6).

Overall, thousands of AREB/ABF-target associations were identified: 14,374 for *A. thaliana*, 10,558 for *S. irio*, 10,197 for *S. parvula*, and 20,026 for *E. salsugineum*. We uncovered highly similar ABRE-like motifs as the most predominant binding site, which is of particular significance as no prior information on TF-DNA interactions were available for *S. irio, S. parvula*, and *E. salsugineum* (Fig. 3A, fig. S6-7, data S1-2 and S6). While the core ACGT sequence motif does not exhibit enrichment across gene regions, ABREs and AREB/ABF binding occurs to a greater extent in the 5’ promoter region (fig. S8, data S6). The top 3,000 sites targeted by each AREB/ABF paralog were highly similar within each species (Fig. 3B). To further investigate these relationships, we swapped the cognate genome for a common noncognate genome (*A. thaliana*) during the pull-down, which we termed swap-DAP; again, we found a large overlap of binding positions between the respective AREB/ABF paralogs to the *A. thaliana* genome. Together these results suggest that AREB/ABF-DNA binding dynamics are highly similar in Brassicaceae and between paralogous AREB/ABFs and that differences in the coding sequence of the AREB/ABF likely play a minor role in determining differences in ABA-dependent gene expression (Fig. 3B, fig. S9).

**Fig. 3.**
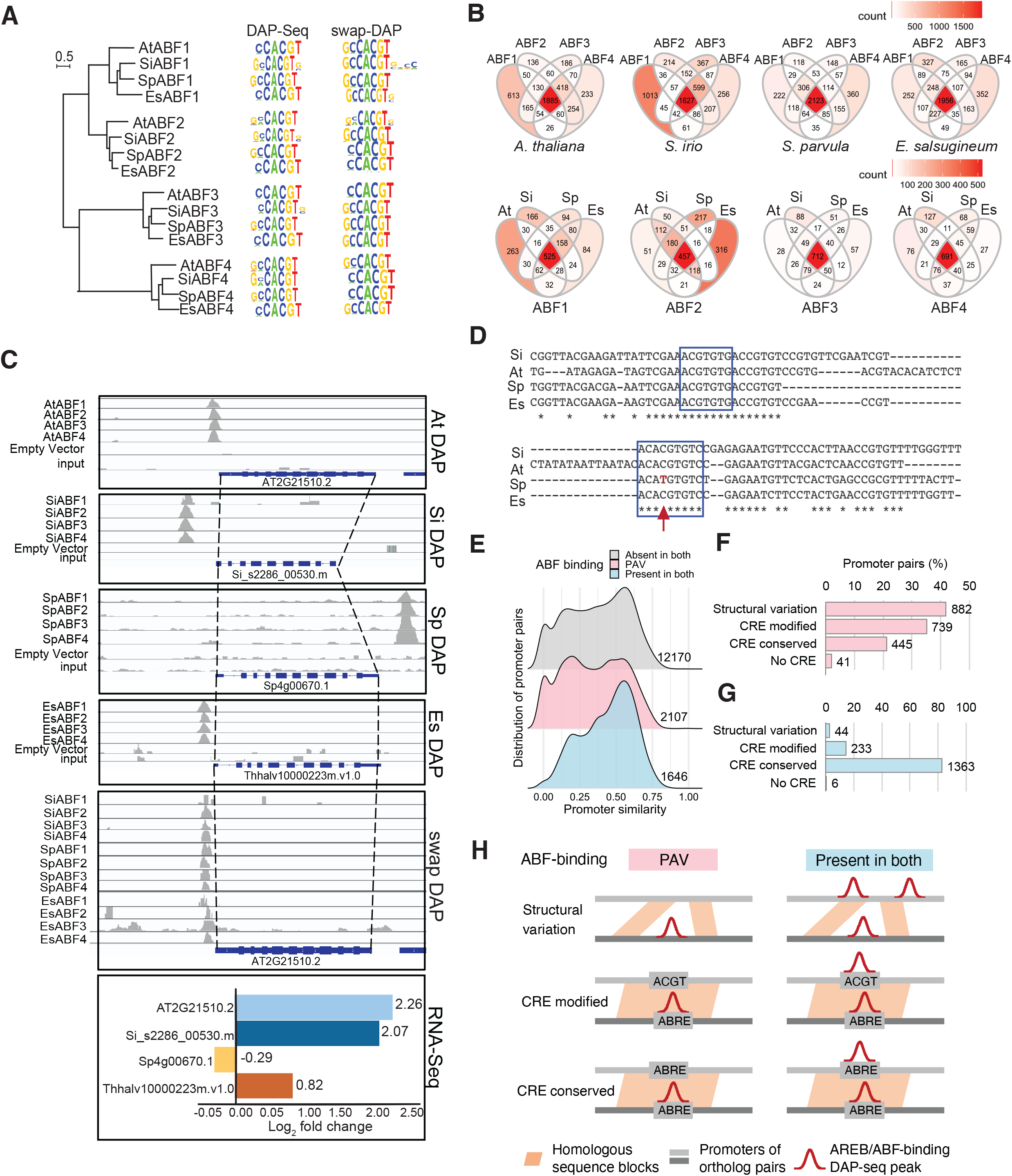
Global ABRE/ABF binding landscape demonstrates dependency between promoter sequence and TF interaction. **(A)** Phylogenetic association of the amino acid sequences of 16 AREB/ABF orthologs and their most enriched motifs. Scale = substitutions/site. **(B)** Venn Diagram showing common peak positions bound by AREB/ABF 1/2/3/4 for top 3000 DAP-Seq peaks. Lower panel shows top 1000 swap-DAP peaks on the *A. thaliana* genome. **(C)** Normalized genome browser window of a locus where AREB/ABF binding is absent exclusively in *S. parvula*, and correlate with gene expression differences. The y-axis for *S. parvula* was adjusted from 10,000 bp to 50 bp scale confirming the absence of peaks. **(D)** A sequence alignment of the 5’ promoters of genes from **(C)**. Blue boxes highlight ABRE motifs. Red arrow indicates the nucleotide change that leads to the loss of the core ABRE motif. **(E-G)** Promoter similarity distribution and sequence variations were compared for 5’ 1Kb promoters of *A. thaliana* and *S. parvula* ortholog pairs that showed conservation or presence/absence variation (PAV) in AREB/ABF binding. Numbers of promoter pairs were shown for each category. **(H)** Mode of sequence variations affecting AREB/ABF binding (see text S1).

We next investigated whether variation in promoter sequence best explained the divergence in ABA response between species. Within our dataset, we observed several incidences where the ABF binding profile deviates in a species-specific manner, an example of which is shown for a molecular chaperone important for stress tolerance (AT2G21510) (Fig. 3C) (*29*). In this case, AREB/ABF binding is associated with greater transcript abundance under 10 µM ABA treatment for all species except *S. parvula*, suggesting that these changes are cis-regulatory effects. Alignment of promoter sequences associated with the AREB/ABF binding site identified a single base pair difference in the core ABRE motif of the *S. parvula* promoter despite other ABRE-like sequences being present (Fig. 3D), suggesting that a well-positioned base pair mutation may be sufficient to disrupt AREB/ABF binding and subsequent gene expression.

To globally establish how differences in AREB/ABF binding relate to promoter sequence variation, we compared the 5’ sequence similarity of orthologous genes, focusing on *A. thaliana* and *S. parvula* (Fig. 3E-H). Gene pairs where both members had significant AREB/ABF binding in this region were much more likely to exhibit sequence similarity across their entire 5’ region compared to gene pairs where neither member, or only one, had an AREB/ABF binding event (Fig. 3E). This pattern of conservation may indicate that additional sequence features, such as other CREs or overall promoter architecture, are needed to facilitate binding of the AREB/ABF. We searched for the presence of ABRE-like sequences within the regions underlying AREB/ABF-binding events deduced from DAP-Seq peaks. For orthologous gene pairs exhibiting presence/absence variation (PAV) in ABF binding, the most frequent underlying difference was the complete gain/loss of the sequence block containing the ABRE (Fig. 3F, “Structural variation”) followed by point mutations disrupting the ABRE (“CRE modified”). Only 21% of PAVs occurred despite a conserved CRE and homologous neighboring sequence, which could be explained by variation in flanking/distal sequences or epigenetic context (*30*). For gene pairs with conserved ABF binding, most promoter pairs had a conserved ABRE sequence underlying the peak (Fig. 3G), as expected. These data highlight the potential role of single nucleotide polymorphisms (SNPs) and insertions/deletions (indels) as frequent drivers of divergence in the ability of promoters to recruit TFs that may lead to the acquisition of lineage specific traits.

We next sought to define an AREB/ABF-centered gene regulatory network (GRN) for ABA signaling that established conserved, core components of the network as well as peripheral modules that exhibited species-specific divergence (*31*). A conserved GRN was established by identifying 354 high confidence AREB/ABF targets (HCATs) that responded across species (320 for 3 hours root, 84 for 24 hours root, 45 for 3 hours shoot, and 47 for 24 hours shoot) and had AREB/ABF binding (Fig. 4A). While AREB/ABF binding to the 5’ proximal region of genes was best correlated with differential gene expression (fig. S10, table S2), we observed that orthologous HCAT genes also showed conserved patterns of AREB/ABF binding in the gene body or 3’ untranslated regions (Fig. 4B, upper panel). The magnitude of ABA regulation for these genes was also highly conserved across species (Fig 4B, lower panel).

**Fig. 4.**
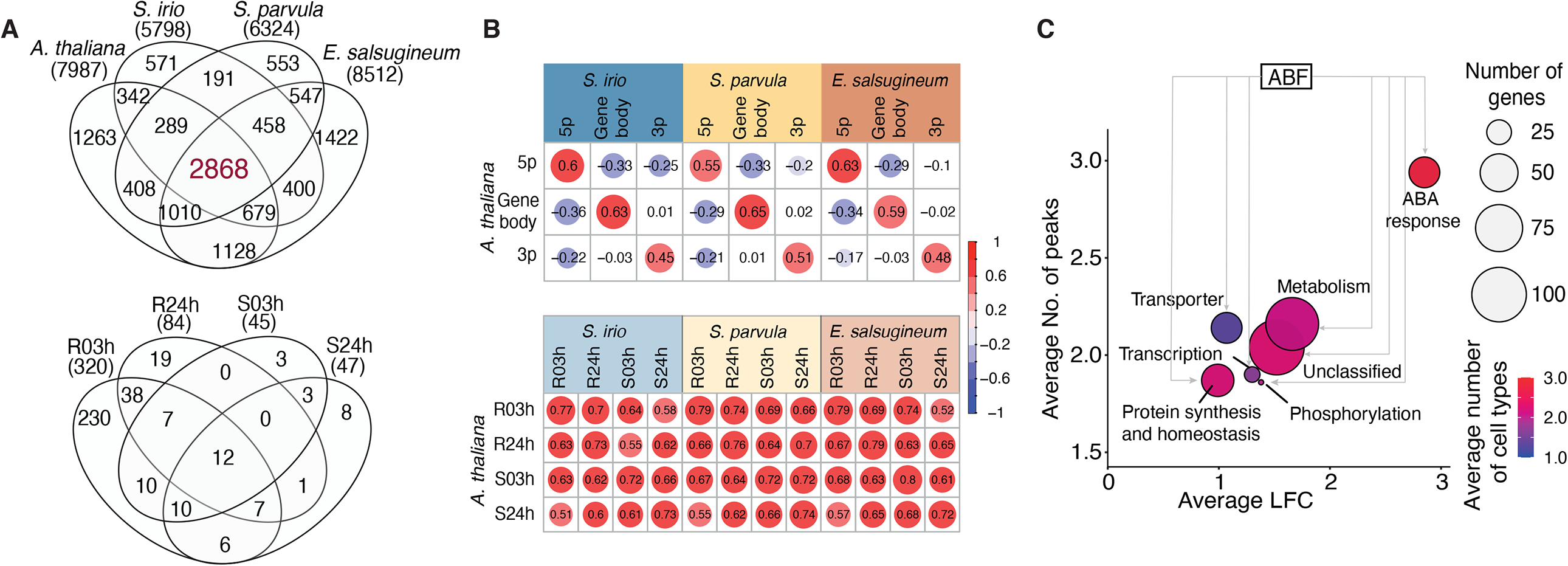
A high confidence ABRE/ABF regulatory network in Brassicaceae. **(A)** Top: Venn diagram of a shared AREB/ABF regulatory module for 1:1 ortholog groups where all orthologs are associated with a DAP-Seq peak (2,898). Bottom: Venn diagram of shared AREB/ABF target genes where all orthologs were differentially expressed (3 hours roots (R03h), 24 hours roots (R24h), 3 hours shoots (S03h), 24 hours shoots (S24h), (*Padj*<0.05). These genes were considered as high confidence ABF targets (HCATs). **(B)** Top: Correlation plot comparing the number of AREB/ABF binding sites (peaks) at different genomic features of HCATs, in *A. thaliana* compared with other species. Bottom: Correlation plot showing the conservation of ABA response, Log_2_ fold change (LFC), in HCATs, circles indicate significant correlations calculated by Spearman’s correlation (ρ, *Padj*<0.05). **(C)** A bubble diagram illustrating the relationship between average LFC (x-axis) and the average number of peaks associated with an ortholog group (y-axis) for HCATs. Bubble size indicates the number of genes, bubble color indicates average NaCl-responsive cell types. “ABA response” genes correlate with greater number of peaks, a broader activation domain upon salt stress, and larger LFC response, while “transporter” showed fewer number of peaks, narrower expression domain, and lower LFC response.

We used these HCATs to construct a core conserved ABF network and identified functional categories that were directly targeted by AREB/ABFs and regulated by ABA across all 4 species (data S7). Curation of the genes in this network revealed that known ABA-specific TFs and signaling components were well represented and tended to have multiple AREB/ABF binding sites in their promoters, which correlated with a greater magnitude of ABA induction across species and broad activation of expression across tissue layers in roots during salt stress, based on data from *A. thaliana* (*32*) (Fig 4C). Our findings also suggest that AREB/ABFs induce more nuanced tissue-specific changes in physiological responses, such as transport, which may be facilitated by having fewer AREB/ABF binding sites (Fig 4C).

We next looked for evidence of subcircuit topologies that differed between species to contextualize AREB/ABF GRN divergence in the Brassicaceae. We focused our analysis on the auxin and ethylene pathways, as these hormones are known to cross-talk with ABA to mediate changes in growth (*33*-*36*). In particular, we were curious to identify divergence in AREB/ABF targets that would illuminate our understanding of the growth promoting response of ABA observed in *S. parvula*. We curated a list of genes with roles in biosynthesis, transport, and signaling for 199 auxin related genes and 72 ethylene related genes (fig. S11-12) (*37*-*47*). Of these genes, AREB/ABFs targeted 81 auxin related genes and 33 ethylene related genes in at least one species. While we observed GRN differences in 18 out of the 33 ethylene-related genes, only 2 were *S. parvula*-specific. In contrast, we observed 15 auxin-related genes with *S. parvula*-specific GRN differences (Fig. 5A). We also observed transcriptional patterns that correlate with these GRN differences providing *in vivo* evidence for the repurposing of an auxin related subnetwork in *S. parvula* (Fig. 5B).

**Fig. 5.**
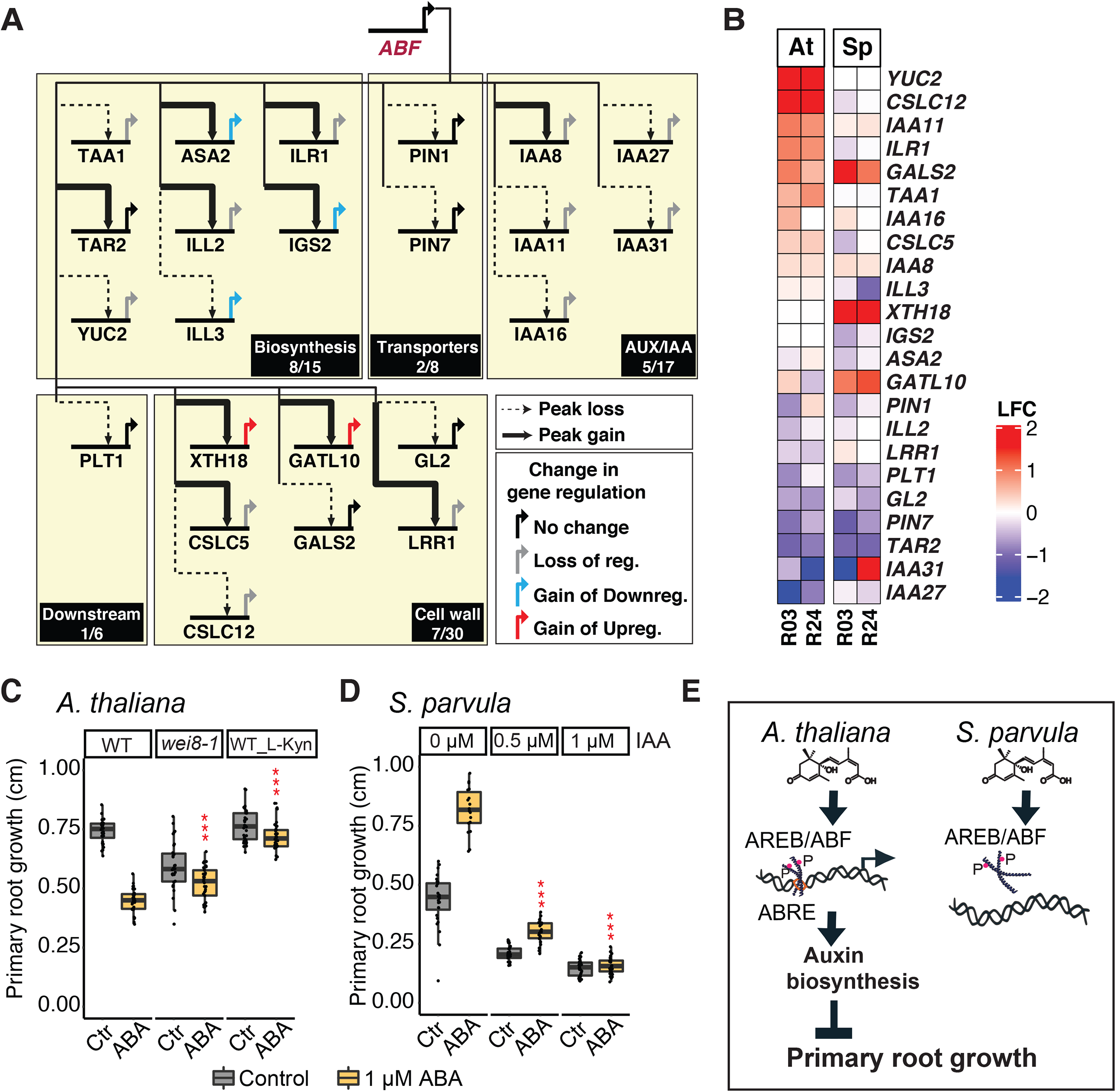
Rewiring of the AREB/ABF-auxin network is responsible for altered growth response in *S. parvula*. **(A)** Divergent GRN for auxin biosynthesis, transport, signaling, and cell wall regulation are highlighted in this network composed based on literature curation. *S. parvula* data was overlaid onto the *A. thaliana* network. Species-specific differences are highlighted as well as the gain/loss of a peak based on AREB/ABF binding. **(B)** A heatmap summarizing the RNA-Seq data across *A. thaliana* and *S. parvula* for species-specific and lineage-specific ABA responses highlighted in the GRN. **(C)** Quantification of primary root growth for wild-type (WT) and *wei8-1* mutants treated with mock, 1 µM ABA, as well as WT with 1µM L-kynurenine (L-Kyn), and 1 µM L-Kyn + 1 µM ABA. Asterisk indicates significance by 2-way ANOVA. \*\*\**P* < 0.001, n>30. **(D)** Quantification of primary root growth for *S. parvula* treated with control, 1 µM ABA, as well as 0.5 µM IAA, 1 µM IAA and co-treatment of IAA and ABA. Asterisk indicates significance by 2-way ANOVA. \*\*\**P* < 0.001, n>25. **(E)** A schematic representation of the mechanism we proposed for the differential regulation in primary root growth in *S. parvula* compared with *A. thaliana*.

Auxin inhibits root growth by suppressing the rate of cell elongation (*48*). Establishing an AREB/ABF-mediated GRN in *S. parvula* revealed a large number of genes that encode enzymes involved in auxin biosynthesis (Fig. 5A). These include *TAA1* and *YUC2*, which mediate the last steps of auxin biosynthesis, as well as root growth inhibition in response to stress (*49, 50*). In *A. thaliana*, AREB/ABF binding likely leads to the induction of these genes upon ABA treatment, while in *S. parvula*, AREB/ABF binding is lost, and their expression is unaffected by ABA treatment. Thus, we hypothesized that differences in the ability of ABA to inhibit growth may, in part, be mediated by differential control of *TAA1* or *YUC2*. We tested our hypothesis by using a TAA1 inhibitor, L-Kyruneine, and the TAA1 mutant allele *wei8-1*, and found that both methods of reducing auxin biosynthesis lowered the sensitivity of roots to ABA treatment in *A. thaliana*, thus partially phenocopying what we observed in *S. parvula* under ABA treatment (*51, 52*). These data are consistent with previous work demonstrating the role of auxin signaling in mediating ABA-response (*33*). Finally, we found that the growth inducing effects of ABA treatment on *S. parvula* could be reversed with supplemental auxin (Fig. 5D), thus recapitulating an *A. thaliana*-like signaling environment. Together our findings reveal that *S. parvula* has undergone extensive rewiring of its AREB/ABF network, with specific changes in the connectivity to the auxin pathway and growth regulation (Fig. 5E).

## Conclusions

We used species within the Brassicaceae to investigate how stress-mediated GRNs differed between extremophyte and stress sensitive species, taking advantage of the extensive gene annotation and knowledge associated with *A. thaliana*. In building an AREB/ABF-centered GRN, we identified a robust evolutionarily conserved ABA pathway, and divergence in peripheral pathways that contribute to hormone cross-talk and growth regulation.

Previous work has proven the utility of manipulating the promoter sequences of plant growth regulators to generate agriculturally beneficial traits in crops (*53*). The work presented here has identified genes whose promoters have likely been the target of natural selection for extreme stress tolerance and will likely fuel the further implementation of a genetic engineering strategy for crop improvement that targets gene regulatory sequences. Future work to incorporate more Brassicaceae species into a comparative framework will further elucidate the principles by which divergence in GRNs contribute to novel physiological outcomes and adaptation to stressful environments.

## Acknowledgments

We would like to acknowledge Mary Galli and Muh Ching Yee for advice on experimental design, Carnegie Institution for Science, Department of Plant Biology and Garret Huntress for providing access to computing resources and data management, undergraduates Rachel Gates and Jen Pulido for their summer engagements with this study, and Chris Pires for *S. irio* seeds.

## Funding

US Department of Energy’s Biological and Environmental Research program (Grant DE-SC0020358, to J.R.D., D.H.O and M.D)

Carnegie Institution for Science endowment (J.R.D)

National Science Foundation (MCB-1616827 and NSF-IOS-EDGE-1923589) (DO and MD) RDA, South Korea (Next-Generation BioGreen21 program PJ01317301) (DO and MD) National Science Foundation Graduate Research Fellowship (Y.S.)

HHMI-Simons Faculty Scholar (J.R.D)

## Author contributions

YS and LD performed the experiments.

YS, DO, PR, AR and JRD analyzed the data.

YS and JRD wrote the manuscript.

DO, LD, and PR contributed to the manuscript preparation.

## Competing interests

Authors declare no competing interests.

## Data and materials availability

All data are available in the manuscript, in the supplementary material, or in the following databases: high-throughput sequencing data sets are available through the National Center for Biotechnology Information Sequence Read Archive (NCBI SRA) under BioProject ID PRJNA682697 (https://www.ncbi.nlm.nih.gov/bioproject/682697). The custom scripts used to analyze the data, along with step-by-step procedures, are available at https://github.com/dinnenylab/BrassicaceaeGRN. Supplementary data are available on FigShare at https://figshare.com/s/988a4f4996a97f7824a1 (DOI:10.6084/m9.figshare.14033822). Genome browser view of data can be found on Jbrowse: http://jbrowsedap.s3-website-us-west-1.amazonaws.com.

## Supplementary Materials

Materials and Methods

Figures 1-5

References (*01-69*)

## Notes

https://github.com/dinnenylab/BrassicaceaeGRN

### Competing Interest Statement

The authors have declared no competing interest.

### Summary of Updates

Updated text and figures

## References and Notes

1. Y. Kazachkova, G. Eshel, P. Pantha, J. M. Cheeseman, M. Dassanayake, S. Barak, Halophytism: what have we learnt from Arabidopsis thaliana relative model systems? Plant Physiol. 178, 972–988 (2018).

2. A. Raza, A. Razzaq, S. S. Mehmood, X. Zou, X. Zhang, Y. Lv, J. Xu, Impact of climate change on crops adaptation and strategies to tackle its outcome: a review. Plants. 8 (2019), doi:10.3390/plants8020034.

3. J.-K. Zhu, The next top models. Cell (2015).

4. A. O. Shamustakimova, T. G. Leonova, V. V. Taranov, A. H. de Boer, A. V. Babakov, Cold stress increases salt tolerance of the extremophytes Eutrema salsugineum (Thellungiella salsuginea) and Eutrema (Thellungiella) botschantzevii. J. Plant Physiol. 208, 128–138 (2017).

5. . A. Mishra, B. Tanna, Halophytes: potential resources for salt stress tolerance genes and promoters. Front. Plant Sci. 8, 829 (2017).

6. H.-J. Wu, Z. Zhang, J.-Y. Wang, D.-H. Oh, M. Dassanayake, B. Liu, Q. Huang, H.-X. Sun, R. Xia, Y. Wu, Y.-N. Wang, Z. Yang, Y. Liu, W. Zhang, H. Zhang, J. Chu, C. Yan, S. Fang, J. Zhang, Y. Wang, F. Zhang, G. Wang, S. Y. Lee, J. M. Cheeseman, B. Yang, B. Li, J. Min, L. Yang, J. Wang, C. Chu, S.-Y. Chen, H. J. Bohnert, J.-K. Zhu, X.-J. Wang, Q. Xie, Insights into salt tolerance from the genome of Thellungiella salsuginea. Proc. Natl. Acad. Sci. U. S. A. 109, 12219–12224 (2012).

7. M. Dassanayake, D.-H. Oh, J. S. Haas, A. Hernandez, H. Hong, S. Ali, D.-J. Yun, R. A. Bressan, J.-K. Zhu, H. J. Bohnert, J. M. Cheeseman, The genome of the extremophile crucifer Thellungiella parvula. Nat. Genet. 43, 913–918 (2011).

8. W. Xuejie, Z. L. F. Shoujin, Comparative studies on growth and physiological reaction of Thellungiella Salsuginea under NaCl and Na2SO4 stresses. Journal of Shandong Normal University. 1 (2007) (available at https://en.cnki.com.cn/Article_en/CJFDTotal-SDZK200701042.htm).

9. Y. P. Lee, C. Funk, A. Erban, J. Kopka, K. I. Köhl, E. Zuther, D. K. Hincha, Salt stress responses in a geographically diverse collection of Eutrema/Thellungiella spp. accessions. Funct. Plant Biol. 43, 590–606 (2016).

10. J. Wang, Q. Zhang, F. Cui, L. Hou, S. Zhao, H. Xia, J. Qiu, T. Li, Y. Zhang, X. Wang, C. Zhao, Genome-wide analysis of gene expression provides new insights into cold responses in Thellungiella salsuginea. Front. Plant Sci. 8, 713 (2017).

11. D.-H. Oh, H. Hong, S. Y. Lee, D.-J. Yun, H. J. Bohnert, M. Dassanayake, Genome structures and transcriptomes signify niche adaptation for the multiple-ion-tolerant extremophyte Schrenkiella parvula. Plant Physiol. 164, 2123–2138 (2014).

12. M. J. R. MacLeod, J. Dedrick, C. Ashton, W. W. L. Sung, M. J. Champigny, E. A. Weretilnyk, Exposure of two Eutrema salsugineum (Thellungiella salsuginea) accessions to water deficits reveals different coping strategies in response to drought. Physiol. Plant. 155, 267–280 (2015).

13. F. Orsini, M. P. D’Urzo, G. Inan, S. Serra, D.-H. Oh, M. V. Mickelbart, F. Consiglio, X. Li, J. C. Jeong, D.-J. Yun, H. J. Bohnert, R. A. Bressan, A. Maggio, A comparative study of salt tolerance parameters in 11 wild relatives of Arabidopsis thaliana. J. Exp. Bot. 61, 3787–3798 (2010).

14. H. Claeys, S. Van Landeghem, M. Dubois, K. Maleux, D. Inzé, What Is stress? Dose-response effects in commonly used in vitro stress assays. Plant Physiology. 165, 519–527 (2014).

15. S. R. Cutler, P. L. Rodriguez, R. R. Finkelstein, S. R. Abrams, Abscisic acid: emergence of a core signaling network. Annu. Rev. Plant Biol. 61, 651–679 (2010).

16. S. K. Sah, K. R. Reddy, J. Li, Abscisic acid and abiotic stress tolerance in crop plants. Front. Plnt. Sci. 7, 571 (2016).

17. K. Vishwakarma, N. Upadhyay, N. Kumar, G. Yadav, J. Singh, R. K. Mishra, V. Kumar, R. Verma, R. G. Upadhyay, M. Pandey, S. Sharma, Abscisic acid signaling and abiotic stress tolerance in plants: A review on current knowledge and future prospects. Front. Plant Sci. 8, 161 (2017).

18. T. Yoshida, Y. Fujita, K. Maruyama, J. Mogami, D. Todaka, K. Shinozaki, K. YamaguchiLshinozaki, Four Arabidopsis AREB / ABF transcription factors function predominantly in gene expression downstream of SnRK2 kinases in abscisic acid signalling in response to osmotic stress. Plant, Cell & Environment. 38, 35–49 (2015).

19. K. Yamaguchi-Shinozaki, K. Shinozaki, A novel cis-acting element in an Arabidopsis gene is involved in responsiveness to drought, low-temperature, or high-salt stress. Plant Cell. 6, 251–264 (1994).

20. D. Deprost, L. Yao, R. Sormani, M. Moreau, G. Leterreux, M. Nicolaï, M. Bedu, C. Robaglia, C. Meyer, The Arabidopsis TOR kinase links plant growth, yield, stress resistance and mRNA translation. EMBO Rep. 8, 864–870 (2007).

21. O. Fornes, J. A. Castro-Mondragon, A. Khan, R. van der Lee, X. Zhang, P. A. Richmond, B. P. Modi, S. Correard, M. Gheorghe, D. Baranašić, W. Santana-Garcia, G. Tan, J. Chèneby, B. Ballester, F. Parcy, A. Sandelin, B. Lenhard, W. W. Wasserman, A. Mathelier, JASPAR 2020: update of the open-access database of transcription factor binding profiles. Nucleic Acids Research (2019), doi:10.1093/nar/gkz1001.

22. Y. Fujita, M. Fujita, R. Satoh, K. Maruyama, M. M. Parvez, M. Seki, K. Hiratsu, M. Ohme-Takagi, K. Shinozaki, K. Yamaguchi-Shinozaki, AREB1 is a transcription activator of novel ABRE-dependent ABA signaling that enhances drought stress tolerance in Arabidopsis. Plant Cell. 17, 3470–3488 (2005).

23. F. Li, F. Mei, Y. Zhang, S. Li, Z. Kang, H. Mao, Genome-wide analysis of the AREB/ABF gene lineage in land plants and functional analysis of TaABF3 in Arabidopsis. BMC Plant Biol. 20, 558 (2020).

24. X. Yue, G. Zhang, Z. Zhao, J. Yue, X. Pu, M. Sui, Y. Zhan, Y. Shi, Z. Wang, G. Meng, Z. Zhao, L. An, A cryophyte transcription factor, CbABF1, confers freezing, and drought tolerance in tobacco. Front. Plant Sci. 10, 699 (2019).

25. J. Liu, J. Chu, C. Ma, Y. Jiang, Y. Ma, J. Xiong, Z.-M. Cheng, Overexpression of an ABA-dependent grapevine bZIP transcription factor, VvABF2, enhances osmotic stress in Arabidopsis. Plant Cell Rep. 38, 587–596 (2019).

26. S. Gao, J. Gao, X. Zhu, Y. Song, Z. Li, G. Ren, X. Zhou, B. Kuai, ABF2, ABF3, and ABF4 promote ABA-mediated chlorophyll degradation and leaf senescence by transcriptional activation of chlorophyll catabolic genes and senescence-associated genes in Arabidopsis. Mol. Plant. 9, 1272– 1285 (2016).

27. H. Choi, J. Hong, J. Ha, J. Kang, S. Y. Kim, ABFs, a family of ABA-responsive element binding factors. J. Biol. Chem. 275, 1723–1730 (2000).

28. R. C. O’Malley, S.-S. C. Huang, L. Song, M. G. Lewsey, A. Bartlett, J. R. Nery, M. Galli, A. Gallavotti, J. R. Ecker, Cistrome and epicistrome features shape the regulatory DNA landscape. Cell. 166, 1598 (2016).

29. Y. Guo, S. Mahony, D. K. Gifford, High resolution genome wide binding event finding and motif discovery reveals transcription factor spatial binding constraints. PLoS Comput. Biol. 8, e1002638 (2012).

30. R. Wu, L. Duan, J. Pruneda-Paz, D.-H. Oh, M. P. Pound, S. A. Kay, J. R. Dinneny, The 6xABRE synthetic promoter enables the spatiotemporal analysis of ABA-mediated transcriptional regulation. Plant Physiol. (2018), doi:10.1104/pp.18.00401.

31. Y. Sun, J. R. Dinneny, Q&A: How do gene regulatory networks control environmental responses in plants? BMC Biol. 16, 38 (2018).

32. J. R. Dinneny, T. A. Long, J. Y. Wang, J. W. Jung, D. Mace, S. Pointer, C. Barron, S. M. Brady, J. Schiefelbein, P. N. Benfey, Cell identity mediates the response of Arabidopsis roots to abiotic stress. Science. 320, 942–945 (2008).

33. J. M. Thole, E. R. Beisner, J. Liu, S. V. Venkova, L. C. Strader, Abscisic acid regulates root elongation through the activities of auxin and ethylene in Arabidopsis thaliana. G3: Genes, Genomes, Genetics. 4, 1259–1274 (2014).

34. W. G. Spollen, M. E. LeNoble, T. D. Samuels, N. Bernstein, R. E. Sharp, Abscisic acid accumulation maintains maize primary root elongation at low water potentials by restricting ethylene production. Plant Physiol. 122, 967–976 (2000).

35. J. H. Rowe, J. F. Topping, J. Liu, K. Lindsey, Abscisic acid regulates root growth under osmotic stress conditions via an interacting hormonal network with cytokinin, ethylene and auxin. New Phytol. 211, 225–239 (2016).

36. H. Qin, L. He, R. Huang, The coordination of ethylene and other hormones in primary root development. Front. Plant Sci. 10, 874 (2019).

37. M. Majda, S. Robert, The role of auxin in cell wall expansion. Int. J. Mol. Sci. 19 (2018), doi:10.3390/ijms19040951.

38. L. C. Strader, G. L. Chen, B. Bartel, Ethylene directs auxin to control root cell expansion. Plant J. 64, 874–884 (2010).

39. J. Li, H.-H. Xu, W.-C. Liu, X.-W. Zhang, Y.-T. Lu, Ethylene inhibits root elongation during alkaline stress through AUXIN1 and associated changes in Auxin accumulation. Plant Physiol. 168, 1777– 1791 (2015).

40. H. Takatsuka, M. Umeda, Hormonal control of cell division and elongation along differentiation trajectories in roots. J. Exp. Bot. 65, 2633–2643 (2014).

41. J.-L. Mao, Z.-Q. Miao, Z. Wang, L.-H. Yu, X.-T. Cai, C.-B. Xiang, Arabidopsis ERF1 mediates cross-talk between ethylene and auxin biosynthesis during primary root elongation by regulating ASA1 expression. PLoS Genet. 12, e1005760 (2016).

42. L. Van den Broeck, M. Dubois, M. Vermeersch, V. Storme, M. Matsui, D. Inzé, From network to phenotype: the dynamic wiring of an Arabidopsis transcriptional network induced by osmotic stress. Mol. Syst. Biol. 13, 961 (2017).

43. Z. Yang, L. Tian, M. Latoszek-Green, D. Brown, K. Wu, Arabidopsis ERF4 is a transcriptional repressor capable of modulating ethylene and abscisic acid responses. Plant Mol. Biol. 58, 585–596 (2005).

44. M. Dubois, A. Skirycz, H. Claeys, K. Maleux, S. Dhondt, S. De Bodt, R. Vanden Bossche, L. De Milde, T. Yoshizumi, M. Matsui, D. Inzé, Ethylene Response Factor 6 acts as a central regulator of leaf growth under water-limiting conditions in Arabidopsis. Plant Physiol. 162, 319–332 (2013).

45. M. Müller, S. Munné-Bosch, Ethylene response factors: a key regulatory hub in hormone and stress signaling. Plant Physiol. 169, 32–41 (2015).

46. M. N. Markakis, T. De Cnodder, M. Lewandowski, D. Simon, A. Boron, D. Balcerowicz, T. Doubbo, L. Taconnat, J.-P. Renou, H. Höfte, J.-P. Verbelen, K. Vissenberg, Identification of genes involved in the ACC-mediated control of root cell elongation in Arabidopsis thaliana. BMC Plant Biol. 12, 208 (2012).

47. Z.-Q. Miao, P.-X. Zhao, J.-L. Mao, L.-H. Yu, Y. Yuan, H. Tang, Z.-B. Liu, C.-B. Xiang, HOMEOBOX PROTEIN 52 mediates the crosstalk between ethylene and auxin signaling during primary root elongation by modulating auxin transport-related gene expression. Plant Cell. 30, 2761–2778 (2018).

48. S. M. Velasquez, E. Barbez, J. Kleine-Vehn, J. M. Estevez, Auxin and cellular elongation. Plant Physiol. 170, 1206–1215 (2016).

49. G. Liu, S. Gao, H. Tian, W. Wu, H. S. Robert, Z. Ding, Local transcriptional control of YUCCA regulates auxin promoted root-growth inhibition in response to aluminium stress in Arabidopsis. PLoS Genet. 12, e1006360 (2016).

50. Q. Wang, G. Qin, M. Cao, R. Chen, Y. He, L. Yang, Z. Zeng, Y. Yu, Y. Gu, W. Xing, W. A. Tao, T. Xu, A phosphorylation-based switch controls TAA1-mediated auxin biosynthesis in plants. Nat. Commun. 11, 679 (2020).

51. D. Pacheco-Villalobos, M. Sankar, K. Ljung, C. S. Hardtke, Disturbed local auxin homeostasis enhances cellular anisotropy and reveals alternative wiring of auxin-ethylene crosstalk in Brachypodium distachyon seminal roots. PLoS Genet. 9, e1003564 (2013).

52. A. N. Stepanova, J. Robertson-Hoyt, J. Yun, L. M. Benavente, D.-Y. Xie, K. Dolezal, A. Schlereth, G. Jürgens, J. M. Alonso, TAA1-mediated auxin biosynthesis is essential for hormone crosstalk and plant development. Cell. 133, 177–191 (2008).

53. D. Rodríguez-Leal, Z. H. Lemmon, J. Man, M. E. Bartlett, Z. B. Lippman, Engineering quantitative trait variation for crop improvement by genome editing. Cell. 171, 470–480.e8 (2017).

54. J. Schindelin, I. Arganda-Carreras, E. Frise, V. Kaynig, M. Longair, T. Pietzsch, S. Preibisch, C. Rueden, S. Saalfeld, B. Schmid, J.-Y. Tinevez, D. J. White, V. Hartenstein, K. Eliceiri, P. Tomancak, A. Cardona, Fiji: an open-source platform for biological-image analysis. Nature Methods. 9, 676–682 (2012).

55. B. Patel, Simple, fast, and efficient cloning of PCR products with TOPO® cloning vectors. BioTechniques. 46, 559 (2009).

56. A. Haudry, A. E. Platts, E. Vello, D. R. Hoen, M. Leclercq, R. J. Williamson, E. Forczek, Z. Joly-Lopez, J. G. Steffen, K. M. Hazzouri, K. Dewar, J. R. Stinchcombe, D. J. Schoen, X. Wang, J. Schmutz, C. D. Town, P. P. Edger, J. C. Pires, K. S. Schumaker, D. E. Jarvis, T. Mandáková, M. A. Lysak, E. van den Bergh, M. E. Schranz, P. M. Harrison, A. M. Moses, T. E. Bureau, S. I. Wright, M. Blanchette, An atlas of over 90,000 conserved noncoding sequences provides insight into crucifer regulatory regions. Nat. Genet. 45, 891–898 (2013).

57. R. M. Waterhouse, M. Seppey, F. A. Simão, M. Manni, P. Ioannidis, G. Klioutchnikov, E. V. Kriventseva, E. M. Zdobnov, BUSCO applications from quality assessments to gene prediction and phylogenomics. Mol. Biol. Evol. 35, 543–548 (2018).

58. M. Pertea, D. Kim, G. M. Pertea, J. T. Leek, S. L. Salzberg, Transcript-level expression analysis of RNA-seq experiments with HISAT, StringTie and Ballgown. Nat. Protoc. 11, 1650–1667 (2016).

59. M. I. Love, W. Huber, S. Anders, Moderated estimation of fold change and dispersion for RNA-seq data with DESeq2. Genome Biol. 15, 550 (2014).

60. D.-H. Oh, M. Dassanayake, Landscape of gene transposition-duplication within the Brassicaceae family. DNA Res. 26, 21–36 (2019).

61. M. Steinegger, J. Söding, MMseqs2 enables sensitive protein sequence searching for the analysis of massive data sets. Nat. Biotechnol. 35, 1026–1028 (2017).

62. S. Maere, K. Heymans, M. Kuiper, BiNGO: a Cytoscape plugin to assess overrepresentation of gene ontology categories in biological networks. Bioinformatics. 21, 3448–3449 (2005).

63. G. Wang, D.-H. Oh, M. Dassanayake, GOMCL: a toolkit to cluster, evaluate, and extract non-redundant associations of Gene Ontology-based functions. BMC Bioinformatics. 21, 139 (2020).

64. R. C. McLeay, T. L. Bailey, Motif Enrichment Analysis: a unified framework and an evaluation on ChIP data. BMC Bioinformatics. 11, 165 (2010).

65. P. Virtanen, R. Gommers, T. E. Oliphant, M. Haberland, T. Reddy, D. Cournapeau, E. Burovski, P. Peterson, W. Weckesser, J. Bright, S. J. van der Walt, M. Brett, J. Wilson, K. J. Millman, N. Mayorov, A. R. J. Nelson, E. Jones, R. Kern, E. Larson, C. J. Carey, İ. Polat, Y. Feng, E. W. Moore, J. VanderPlas, D. Laxalde, J. Perktold, R. Cimrman, I. Henriksen, E. A. Quintero, C. R. Harris, A. M. Archibald, A. H. Ribeiro, F. Pedregosa, P. van Mulbregt, SciPy 1.0 Contributors, SciPy 1.0: fundamental algorithms for scientific computing in Python. Nature Methods. 17, 261–272 (2020).

66. D. M. Emms, S. Kelly, OrthoFinder: solving fundamental biases in whole genome comparisons dramatically improves orthogroup inference accuracy. Genome Biol. 16, 157 (2015).

67. B. Langmead, S. L. Salzberg, Fast gapped-read alignment with Bowtie 2. Nature Methods (2012).

68. A. R. Quinlan, I. M. Hall, BEDTools: a flexible suite of utilities for comparing genomic features. Bioinformatics. 26, 841–842 (2010).

69. W. J. R. Longabaugh, BioTapestry: a tool to visualize the dynamic properties of gene regulatory networks. Methods Mol. Biol. 786, 359–394 (2012).

