## Supplemental figures for "Divergence in a stress regulatory network underlies differential growth control"

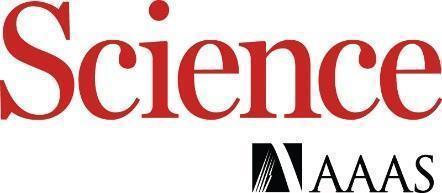

Supplementary Materials for

**Divergence in a stress regulatory network underlies differential growth control**

Ying Sun^1^†, Dong-Ha Oh^2^†, Lina Duan^1^, Prashanth Ramachandran^1^, Andrea Ramirez^1^, Anna Bartlett^3^, Maheshi Dassanayake^2^, José R. Dinneny^1^*

**This PDF file includes:**

Materials and Methods

Supplementary Text

Figs. S1 to S12

Tables S1 to S2

Captions for Data S1 to S8

**Other Supplementary Materials for this manuscript include the following:**

Data S1 to S8

data S1. All 1-to-1 Orthologous groups with functional genomics data from RNA-seq and DAP-seq.

data S2. An analysis with OrthNet (including duplicates) with functional genomics data from RNA-seq and DAP-seq.

data S3. GO enrichment among ABA-induced and repressed DEGs.

data S4. Phylogenetically informed profiling (PiP) analysis of ABA-responsive gene expression for the four crucifer species.

data S5. JASPAR motifs enriched among promoters of ABA-induced and repressed DEGs.

data S6. All DAP-seq peak coordinates with annotation.

data S7. Conserved ABA GRN.

data S8. Auxin and Ethylene GRN.

**Materials and Methods**

Plant Material

*Arabidopsis thaliana* Col-0 ecotype was used in this study. *Sisymbrium irio* was obtained from Chris Pires, University of Missouri (KM 88-34-20-14). *Eutrema salsugineum* ecotype Shandong and *Schrenkiella parvula* ecotype Lake Tuz were from *Arabidopsis* Biological Resource Center (CS22504 and CS22663, respectively).

Phenotypic analysis of primary root growth

Seedling growth and transfer conditions are described in supplementary Materials and Methods. After seedlings are transferred to media containing various supplements, images of seedlings were captured using a CanonScan 9000F flatbed scanner (Canon). Images were quantified using Fiji (*54*). The scale was set as 237 pixels/cm and the image was measured using the segmented line tool. Data visualization and statistical tests were performed using R.

DAP-Seq library preparation and high-throughput sequencing

The coding sequence of the AREB/ABF orthologs were identified using reciprocal blast and defined regions of synteny across species. Gateway cloning compatible primers were designed to amplify each coding sequence from cDNA and clone the coding sequence into TOPO entry vectors (*55*). We found that using rabbit reticulocyte (TnT® T7 Coupled Reticulocyte Lysate System, Recombinant RNasin® Ribonuclease Inhibitor- Promega) instead of wheat germ extract for in vitro transcription and translation of the AREB/ABFs significantly improved the quality of binding events detected. Shoot tissues from 6-day old seedlings were used to generate the genomic DNA library. gDNA was extracted by grinding tissues with cold mortar and pestle then using DNeasy Plant Maxi Kit (Qiagen). All other steps were performed as previously described in (*28*).

Genomes and the use of genomes for bioinformatic analysis

The genome and gene models for *A. thaliana* were obtained from TAIR (Araport11), while genomes and gene models for *S. parvula* (version 2.2) and *E. salsugineum* (version 1.0) were from Phytozome (Phytozome ID 574 and 173, respectively). For *S. irio*, we updated the previously published gene models (*56*) with an updated version using evidence-assisted *ab initio* gene model prediction by the MAKER v. 2.31.10 (https://www.yandell-lab.org/software/maker.html) based on *S. irio* BUSCO-trained parameters (*57*) and RNA-seq data from the current study (fig. S1).

RNA-Seq data analysis

Raw reads were filtered and trimmed using Trim Galore (https://www.bioinformatics.babraham.ac.uk/projects/trim_galore/) which also generated FASTQ files for each of the libraries. Filtered reads were aligned to the reference genome using HISAT2 v. 2.2 and the expression of each primary gene model was estimated with StringTie v. 2.0.1 with default parameters (*58*). Differentially expressed genes (DEGs) were estimated using DESeq2 (*59*) by contrasting ABA-treated samples to controls for each tissue and time point. For comparative analyses, primary protein-coding gene models from the four species were compared using CLfinder pipeline with MMSeqs2 as the aligner, to identify ortholog pairs and groups between pairs of species as well as for all four species (*60*) (*61*). Gene ontology (GO) and known transcription factor binding sites enriched among DEGs and other gene subsets were identified and processed using BiNGO, GOMCL, JASPAR database, and AME and combined into matrices using custom scripts (*62*) (*63*) (*21*) (*64*).

Phylogenetically informed Profiling (PiP) applies Spearman’s rank correlation to the fold-change of ABA responses of orthologs annotated with GO terms between all species pairs. The significance of positive correlation was then estimated for all species pairs using scipy.stats.spearmanr and corrected for multiple testing (Benjamini-Hochberg) for all GO terms and tissue-treatments (*65*). The resulting matrices of presence or absence of significant positive correlation in ABA responses were superimposed to the species tree to identify GO terms and tissue-treatment that show lineage(s)-specific modification patterns in gene regulation. We excluded genes that failed to pass the Cook’s distance test by DESeq2, genes showing lower than 2-fold changes with adjusted *P* value>0.05 (DESeq2), and genes with no DEG among their ortholog pair partners in all species (*59*). The species tree was generated based on all protein-coding gene models using the OrthoFinder pipeline (version 2.2.7) (*66*). GO annotation for *A. thaliana* was obtained from the GO consortium (http://geneontology.org/) and applied to their orthologs showing protein sequence alignment (e<10^-5^) over 70% of total length.

DAP-Seq data analysis

At least 5 million reads per sample were filtered to remove adaptors, low-quality reads, and reads mapping to multiple locations. Bowtie (version 2.3) was used to align reads to their respective genomes (*67*). GEM version 3.4 was used for peak calling after normalizing libraries to both input and empty vector controls (*29*). We defined high confidence AREB/ABF-binding sites as DAP-seq peaks that were present in a minimum of two biological replicates. High confidence peaks were merged, counted, and characterized based on position information such as the occurrence of peaks within the 1bp - 1kb (proximal), 1001 - 2Kb (distal) region of the 5’ and 3’ side of each primary protein-coding gene model. Bedtools merge was used to merge peaks with a maximum distance of 3 nucleotides between the center position across all replicates and AREB/ABFs (*68*). To determine the frequency of overlapping peak positions across AREB/ABFs (e.g. in DAP and swap-DAP experiments) we used 2-3 replicates with comparable library size and ranked all the peaks by their q-value (sum of -log_10_) and normalized peak density as defined by GEM across all selected replicates. For the analysis on the mode of presence/absence variation of DAP-seq peaks, 5’ 1kb promoter sequences between ortholog pairs were aligned using lastz and the presence of *cis*-regulatory elements (CRE) were identified as described previously (*60*) (*30*).

GRN curation and construction

A list of genes involved in the perception, biosynthesis, and signaling of ABA, auxin, and ethylene was manually curated through literature review. Genes involved in the control of cell elongation and root growth were also curated for auxin and ethylene. Among these genes, only those with orthologs in the 4 species were considered for further analysis. BioTapestry was used to build GRNs which consisted of genes with significant fold change in response to ABA in at least one of the four species, summarized in data S1 (*69*). Differences in other species were highlighted relative to *A. thaliana*. For example, a locus with AREB/ABF binding site identified in *A. thaliana* but not *S. parvula* was considered to have an absence of AREB/ABF binding in *S. parvula*, represented by dotted lines. The absence of AREB/ABF binding to *A. thaliana*, while present in *S. parvula* was considered to be a presence in AREB/ABF binding (thick lines). Differences in the number of DAP-seq peaks in the promoter of the target gene were not considered for drawing the edges of the GRN.

Plant growth conditions

All seedlings were grown at constant temperature 24°C with light conditions 14 h light and 10 h dark at 130 µmol m^-2^ s^-1^ light intensity. Seeds were surface sterilized using a 95% ethanol solution followed by a 5-min wash in a 20% bleach/ 0.1% Tween-20 solution, then rinsed with sterile deionized water. Stratification was done at 4°C in the dark for 7 days. “Standard media” is sterilized 0.7% Gelzan media containing ¼ x MS nutrients, 1% sucrose, and 0.05% MES was adjusted to pH 5.7 with 1M KOH. Seeds were grown for 6 days before being transferred to standard media supplemented with NaCl or ABA. For root growth assays, 20-25 seeds were placed in a row across the plate with a 1000 mL pipette tip. After 6 days of growth under standard media conditions, the seedlings were transferred either to standard media plates again or to plates with supplements. The position of the root tip was marked at the time of transfer of the root to distinguish the development of the root before transfer and after transfer.

Plant Material for NGS

Tissues collected for NGS datasets were grown on sterilized mesh (about 10 x 10 cm squares) on the corresponding media plates. 3 rows of seeds (~100 seeds) were plated using a 1000 mL pipette tip. The entire mesh was then transferred using forceps to the corresponding media plate after 6 days. Tissues were separated at the root/shoot junction with a sterile razor blade.

RNA-Seq library preparation and high-throughput sequencing

Total RNA was extracted from the root and shoot tissue using RNeasy plant mini kit (Qiagen 74904) according to the manufacturer’s instructions for three biological replicates collected on separate days. RNA quantity was checked by Qubit (Q32852 Qubit® RNA Assay Kit). RNA quality of each sample was assessed with a 2100 Bioanalyzer (Agilent) and fragment analyzer. RNA-Seq libraries were made using NuGen Universal Plus mRNASeq according to the manufacturer’s instructions. Samples were sequenced on Illumina NextSeq (single-read 75-bp run) with the multiplexed samples described above on 3 lanes.

DAP-Seq library preparation and high-throughput sequencing

Primers were designed to amplify the CDS from different species. TM calculator was used to check that the TM is under 72°C and within 1-2°C apart from each other. CACC was added to the 5’ end of the forward primer to make the PCR product compatible with gateway cloning. The reverse complement of the reverse primer was checked to make sure it is not complementing with overhanging sequence GTGG at the 5’end. To make sure that the fused product is going to be in-frame, plasmid maps were generated and the sequence from the ATG of the protein to VP16 was translated using Expasy protein translation. Primers were ordered from IDT and diluted to 100µM with sterile water prior to use.

After the successful construction of entry vectors with AREB/ABF coding sequences, colony PCR and sequencing were done to validate all cloned sequences. The confirmed entry clones were then recombined into a destination vector with LR clonase to generate a vector with an N-terminal HaloTag that is translationally fused to AREB/ABF. The HaloTag is a mutated hydrolase that covalently binds to Halolink resin which allows for stringent washing and the generation of an affinity matrix with the transcription factor. To confirm the successful expression and synthesis of each AREB/ABF, an immunoblot with anti-halo antibody (Anti-HaloTag® Monoclonal Antibody-Promega) was performed to confirm protein expression and binding efficiency for each TF. This ensures at least 50 ng of protein was used for each DAP-Seq pull down.

 	The quality of the TF-affinity matrix was assayed by using western blots to quantify the abundance of the TF before and after binding to the affinity matrix. GST-Halo was used as a protein standard (Promega). 1 µL of reticulocyte extract before binding with beads and 2 µL of extract after the binding was used for each western on a precast gel (4–20% Criterion™ TGX™ Precast Midi Protein Gel, 12+2 well, 45 µl, BioRad). Before loading, Samples were mixed with Lamneii buffer and heated for 5 min at 95°C. Samples were run at 20 mA for 30 min then at 100 V until the bottom was reached. 5x Running buffer is made using Tris 15.1g, Glycine 94g, 50 mL 10% SDS. After running, proteins were transferred onto the nitrocellulose membrane using a midrange, semi-dry system (invitrogen). 10X TBS- 1L (80g NaCl, 2g KCL, 30g Tris base, pH 8 with HCL autoclave for 30 min was diluted to 1X TBST with 0.05% Tween-20 and sodium azide. This was mixed with milk powder to make a 5% blocking buffer. The membrane was blocked for at least 1 hour at room temperature or overnight at 4°C. After 152 blocking, 10µL of anti-halo was added to the 15mL blocking buffer (10µg) for primary antibody incubation. After incubation, the membrane was washed 5x for 5 min each in 1x TBST. The membrane was then incubated for 1 hour with secondary anti-mouse- HRP 1:10,000 in 5% milk-TBST (2µL in 20 mL blocking buffer). The membrane was washed again 5x for 5 min each in 1x TBST. Chemiluminescent reagent in 1:1 ratio was added to the membrane afterward and Chemi (ECL) signal was detected using Sapphire biomolecular imager. Image processing and labeling was done in Fiji.

 	After gDNA extraction from shoot tissue, gDNA was concentrated using ethanol precipitation. gDNA was incubated for 10-15 min with 10% NaOAC, 200% cold 100% ethanol. After precipitation, the sample was washed with 80% ethanol then dried into a pellet, then redissolved. gDNA was fractionated using the COVARIS system under the settings (mode: Frequency sweeping, Duty cycle: 10%, intensity: 5, Cycles burst: 200, Time: 60 seconds, and number of cycles: 3). Another round of ethanol precipitation was done in the same manner as above. End repair (End-it kit; epicentre cat # ER0720 or ER81050- we used this one) and A-tailing (100mM dATP; bioPioneer inc, Klenow; (3'- 5' exo- NEB) # M0212L (1,000 units) (5,000 U/mL) was done prior to ligating fractionated gDNA with partial next-generation sequencing adaptors using Y adaptors. Y adaptors were made by combining Adapter A with Adapter B then incubating the oligo mixture to 86°C for 2 minutes before allowing the reaction to return to room temperature.

 	Quality of DAP library was assayed using qPCR. 2 µL of (5 ng/µL) DNA template was added to 18 µL master mix made with 4µL Phusion high Fidelity DNA polymerase (M0530S, ThermoFisher), 1 µL of 10 mM dNTP, 0.4 µL of 10mM Illumina TruSeq Universal primer and TruSeq Index primer, Phusion enzyme, 10x SYBR Green I (diluted in DMSO), and H2O. DNA without adaptor ligation and water is used as a negative control. Each sample was repeated 3x and on 384 qPCR plates. Thermocycling settings were 2 min at 95°C, 30s at 98°C, 30 cycles of (15s at 98°C, 30 s at 60°C, 1 min at 72°C), followed by hold at 4°C. Quality of the library was assessed by looking at the amplification curve by plotting the curves in excel as CT vs relative fluorescence unit (RFU). If the curve plateaus within 10 cycles, then the library is good. However, if the curve plateaus after 20 cycles, then the library will not work in the final sequencing.

 	Final DNA is quantified using Qubit (broad range) and incubated with the AREB/ABF affinity matrix (Magne HaloTag beads, 20% slurry, promega, PBS, Ph 7.4 (homemade): Dissolve 8g NaCl, 0.2g KCL, 1.44g Na2HPO4 and 0.24g KH2PO4 in 800 mL sterile water. Adjust pH to 7.4 with HCL. Add water to 1 liter and sterilize by autoclaving. Add 200µL of 25% NP40 and mix well, Magnetic rack, DYNAL magnetic bead separations, Invitrogen) to pull down genomic DNA that bind to each AREB/ABF. Full adaptors containing unique 6 bp barcodes were ligated to each eluted fraction of gDNA and multiplexed for high-throughput Next-generation sequencing. Samples were sequenced on Illumina Hi-Seq (paired-read 150-bp run) with the multiplexed samples.

**Supplementary Text**

text S1. Presence and absence variations (PAVs) of AREB/ABF-binding between promoters of ortholog pairs

For all 1:1 ortholog pairs, we explored the sequence variations in the 5’ 1Kb promoter regions and their effects on AREB/ABF-binding. Promoter pairs were categorized as an AREB/ABF-binding DAP-Sea peak “present in both” (Fig. 3E, blue), only in one of the pairs which we termed “presence/absence variation (PAV)” (Fig. 3E, pink), or absent in both (Fig. 3E, grey). We identified homologous sequence blocks based on pairwise sequence alignment using lastz (e <10^-3^). Promoter similarity was defined as the proportion covered with homologous sequence blocks multiplied by percent identity, for 5’ 1Kb promoter sequence of a 1:1 ortholog pair. We found that promoter pairs with an AREB/ABF-binding DAP-Seq peak present in both tend to have higher promoter similarity compared to promoter pairs with AREB/ABF-binding PAV or AREB/ABF-biding absent in both. While we showed the distribution of promoter similarity for *A. thaliana* - *S. parvula* ortholog pairs in Fig. 3E, we observed similar patterns for all species pairs.

For promoter pairs with AREB/ABF-binding present in both and PAV, we compared the occurrences of different types of promoter sequence variations (Fig. 3F-H). If a promoter pair does not have any detectable homologous sequence block or the AREB/ABF-binding DAP-seq peak does not coincide with a homologous sequence block, the promoter pair is considered to include AREB/ABF-binding events associated with a structural variation. For promoter pairs with PAV, if a promoter contains an AREB/ABF DAP-seq peak coinciding with a putative *cis*-regulatory element (CRE), either an ABRE or an ACGT core, and included in a homologous sequence block, the promoter pair was classified depending on whether the other promoter have a corresponding CRE within the homologous sequence block (“CRE conserved”) or not (“CRE modified”) (Fig. 3H). For promoter pairs in the “present in both” group, the same CREs need to be found coinciding with an AREB/ABF DAP-seq peak in both promoters to be considered as “CRE conserved.” Rare occasions where AREB/ABF DAP-seq peak appears in a homologous promoter block without ABRE or ACGT were classified as “No CRE”.

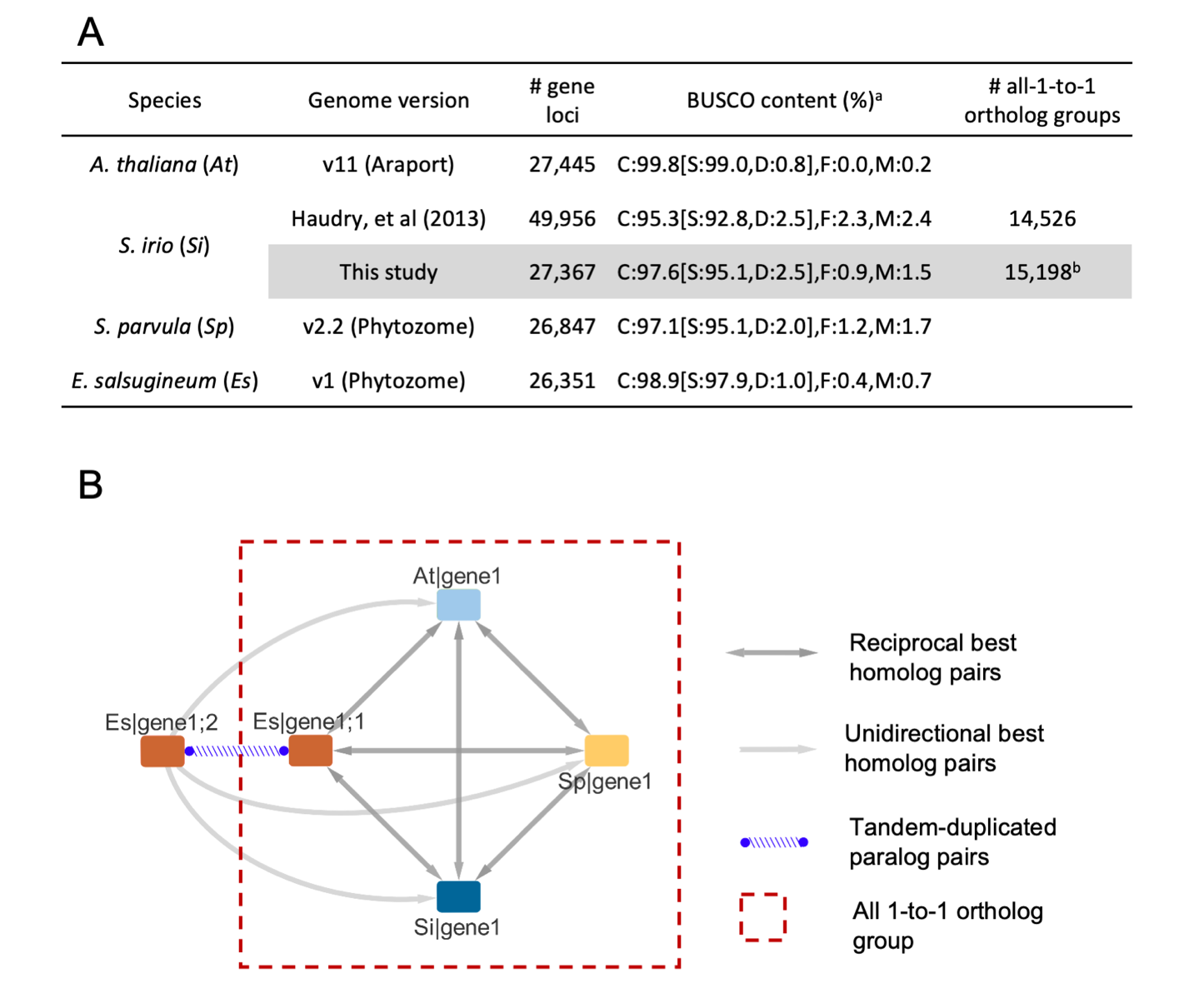

**fig. S1. Genome version and detection of 1-to-1 ortholog groups across species.**

(**A**) The table shows the genome version used for the study as well as gene loci, percent of loci in large scaffolds, and Benchmarking Universal Single-Copy Orthologs (BUSCOs) scores. Gene model annotation for *Sisymbrium irio* was updated for this study. **(B)** OrthNet was constructed to identify the best homologs in other species using protein-coding gene models. Within each OrthNet, a quartet of orthologs unambiguously found each other as reciprocal best homologs (red dashed box). We identified 15,198 “all 1-to-1” ortholog groups (OG) among the four crucifers and used them as the framework for cross-species comparisons (data S1-2).

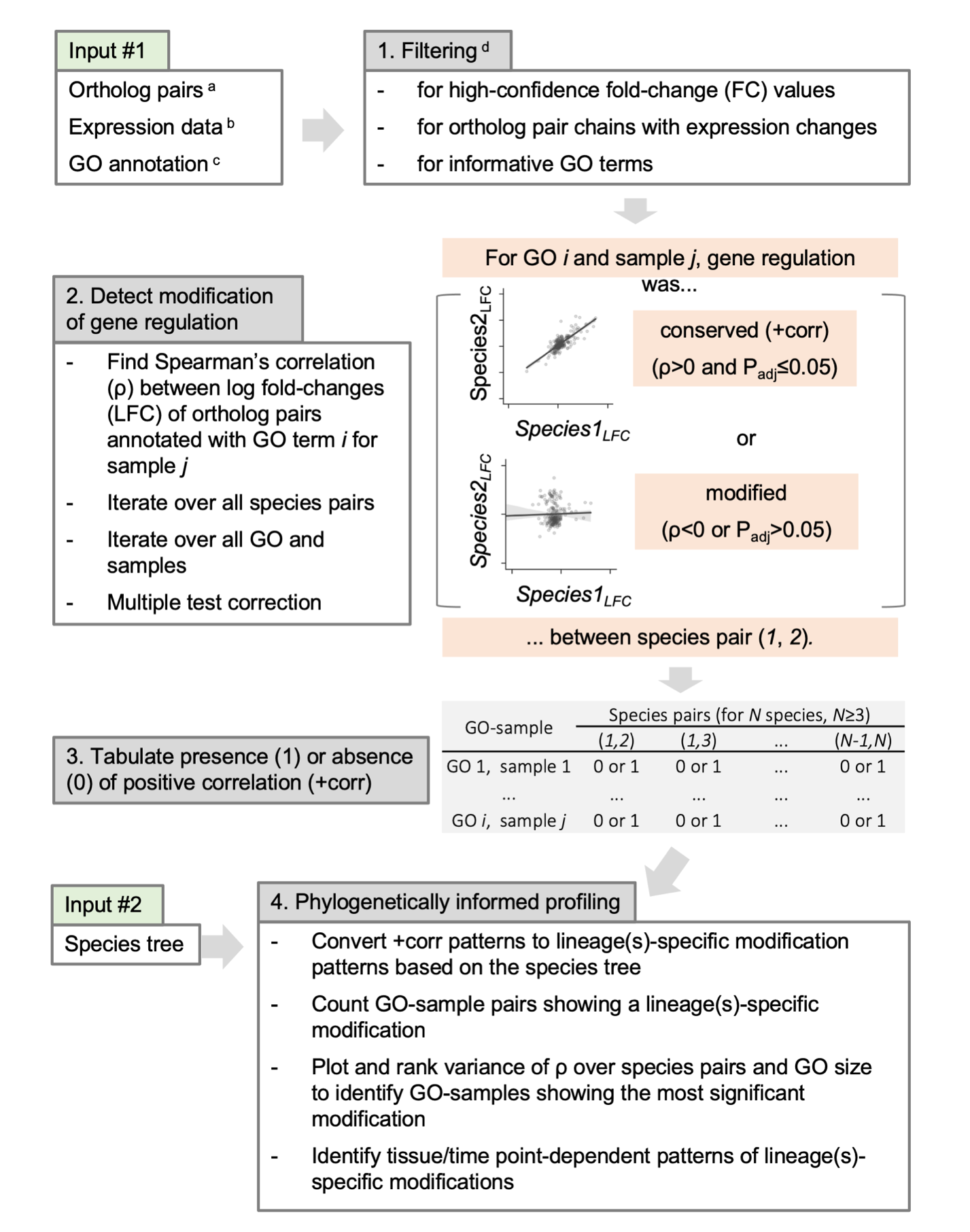

**fig. S2. Overview of the Phylogenetically informed profiling (PiP) pipeline.**

PiP analysis aims to identify GO terms that show lineage(s)-specific modification patterns among multiple closely related species where a majority of genes can be identified as ortholog pairs. Correlations between gene expression patterns of ortholog pairs annotated with each GO term are determined for all species pairs. Presence or absence of significant positive correlations (+corr) are recorded for all species pairs, GO terms, and RNA-seq samples. The +corr patterns are superimposed to the species tree to identify GO terms showing lineage(s)-specific modifications of gene expression in each RNA-seq sample. For the present study: ^a^ For each of all possible species pairs among the four crucifers, best reciprocal homologs are used as ortholog pairs. ^b^ RNA-seq for root and shoot tissues, treated with ABA for 3 or 24 hours. ^c^ For non-Arabidopsis species, a gene coding for a protein with minimum 70% of its length covered by sequence alignment (e<10^-5^) with its best *A. thaliana* homolog was assumed to have the same GO annotation. ^d^ fold-change (FC) was estimated by DESeq2. We removed ortholog pairs including a gene either with non-agreeing RNA-seq replicates (based on using Cook’s distance filter implemented in DESeq2) or with fold-change greater than 2-fold while not a DEG. We removed ortholog pair chains that did not include a DEG in all species pairs. We removed too generic GO terms (i.e. GO size in *A. thaliana* > 2500), redundant GO terms (Jaccard coefficient > 0.8, determined by GOmcl), and GO terms containing less than 10 ortholog pairs in any species pair after filtering.

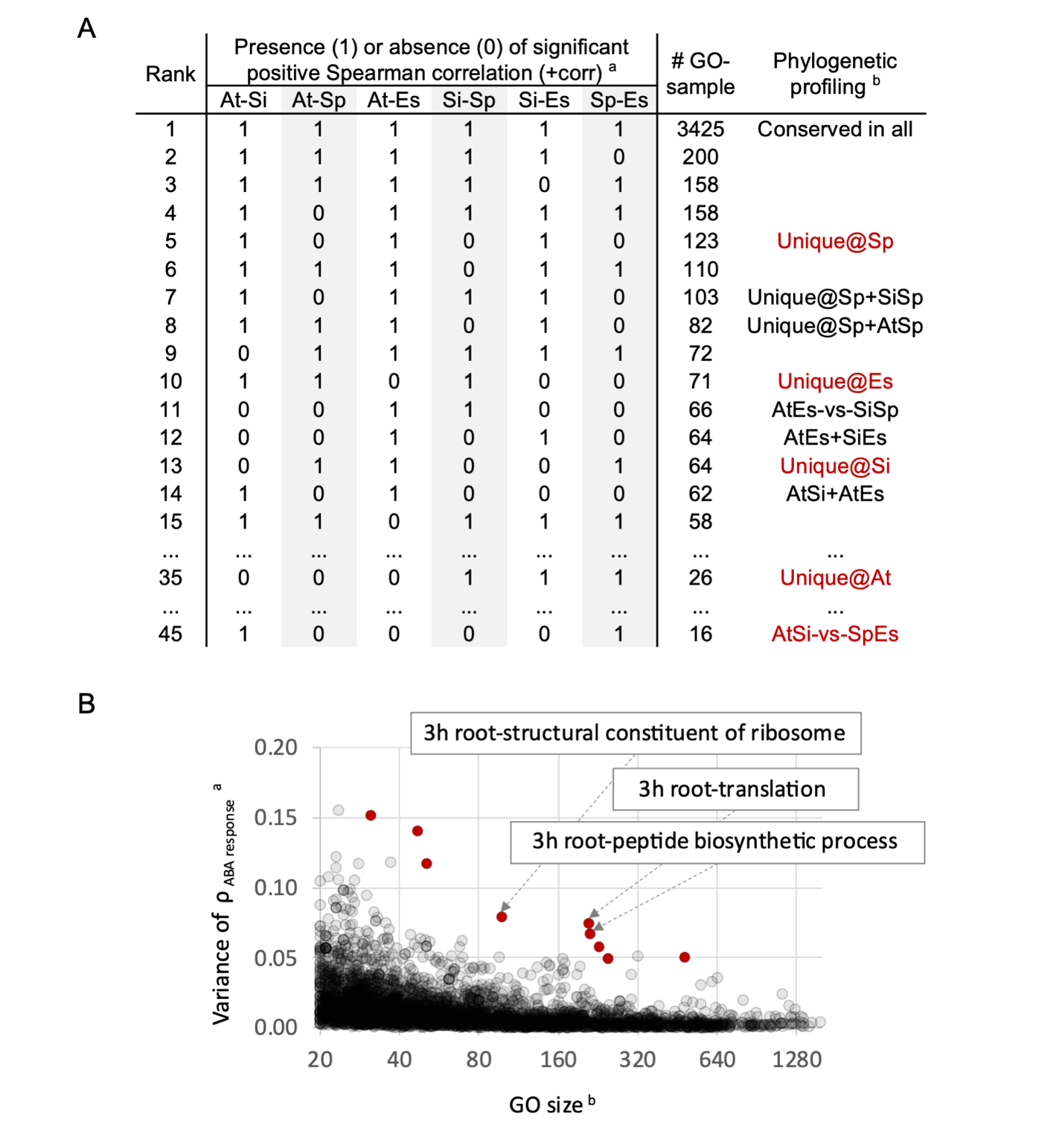

**fig. S3. Phylogenetically informed Profiling (PiP) analysis of ABA-responses.**

**(A)** All non-redundant GO terms were used to determine if there is a significant positive Spearman correlation (+corr) of ABA responses amongst ortholog pairs for all 6 pairs of the 4 species. ^a^ Presence (1) and absence (0) of a significant positive Spearman correlation between ABA-responsive log fold changes at each sample of ortholog pairs annotated with each GO term was shown for all species pairs. ^b^ The most frequent +corr pattern (3,425 GO-sample pairs) represented GO-sample pairs with no modification in any lineage (“Conserved in all”). The next dominant +corr pattern (123 GO-sample pairs) that matches a phylogenetic profile showed a modification of ABA responses uniquely in *S. parvula* (“Unique@Sp”). Phylogenetic profiles “AtEs-vs-SiSp” mirrors the divergence of *S. irio* and *S. parvula* clades from the other two species, while “AtSi-vs-SpEs” indicates potential division between glycophytes (*A. thaliana* and *S. irio*) and halophytes (*S. parvula* and *E. salsugineum*). For phylogenetic profiles marked in red, example GO terms were shown in table S1. **(B)** For all GO-sample pairs, we plotted the variance of Spearman correlation coefficients (ρ) among the six species pairs against the GO size, i.e. the number of ortholog pairs annotated with the GO term and used for PiP analyses. While in general the magnitude of variance was greater for smaller GO terms, “peptide biosynthetic process” and related GO terms in root samples (red) were outliers with greater variances among larger GO terms. Results of PiP analysis for all GO-sample pairs are in data S4.

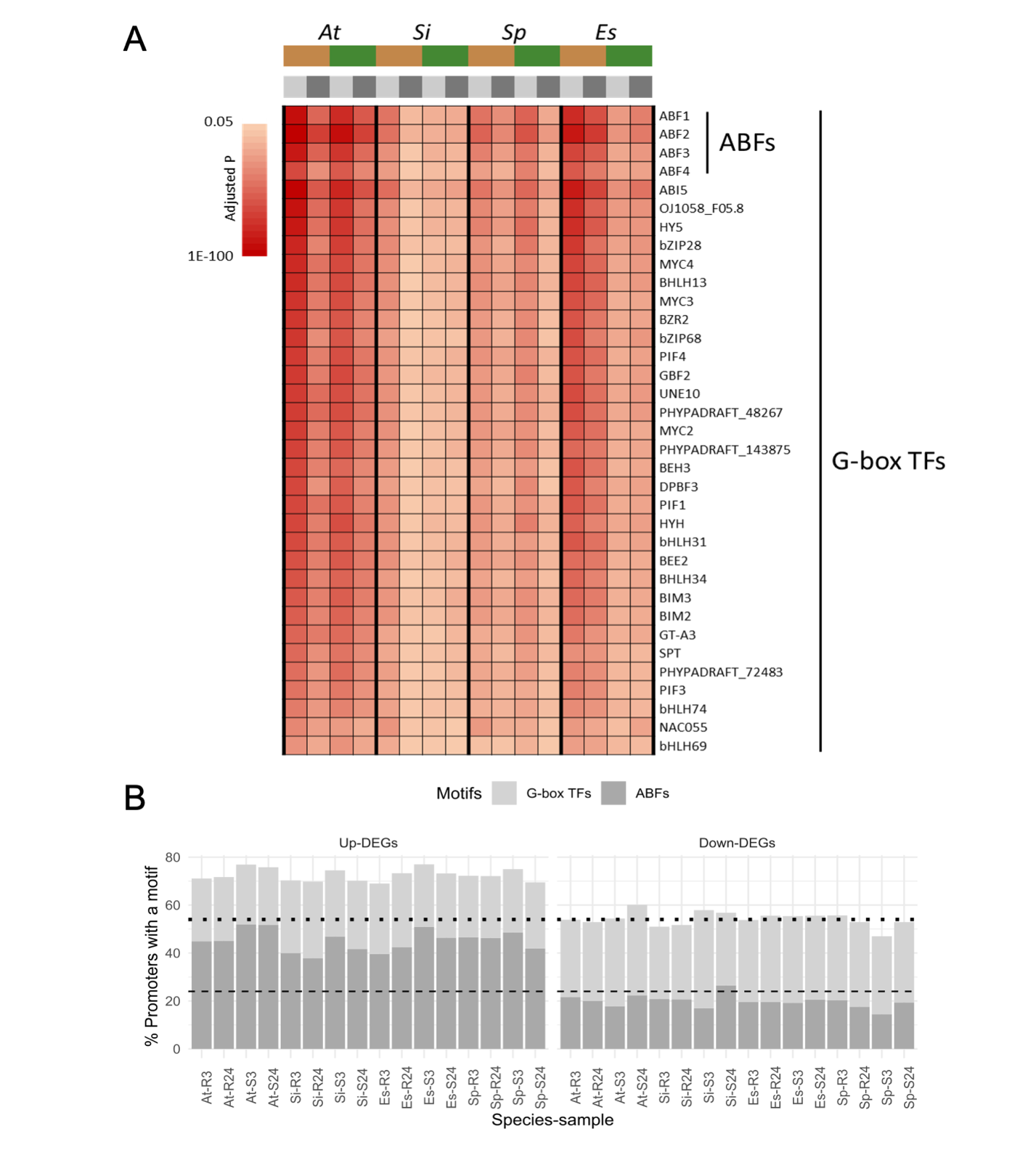

**fig. S4. Transcription factor (TF)-binding motifs enriched among ABA-induced DEGs.**

**(A)** For the 1Kb upstream regions of all ABA-induced and repressed DEGs, we searched for enrichment of known TF-binding motifs (from JASPAR database). The four AREB/ABFs and 31 other G-box-binding TFs (G-box TFs) were the only TFs whose known binding motifs showed significant enrichment (determined using AME) in promoters of ABA-induced DEG promoters in all samples. **(B)** Proportion of promoters including an AREB/ABF or G-box TF-binding motif for both DEGs ABA-induced (Up-DEGs) and repressed (Down-DEGs), with the mean proportion among non-DEGS shown as dotted lines (G-Box TF) and dashed lines (AREB/ABFs). Tissue types and time points are indicated as R3 for 3 hours roots, R24 for 24 hours roots, S3 for 3 hours shoots, and S24 for 24 hours shoots. The enrichment profiles of all JASPAR TF-binding motifs are in data S5.

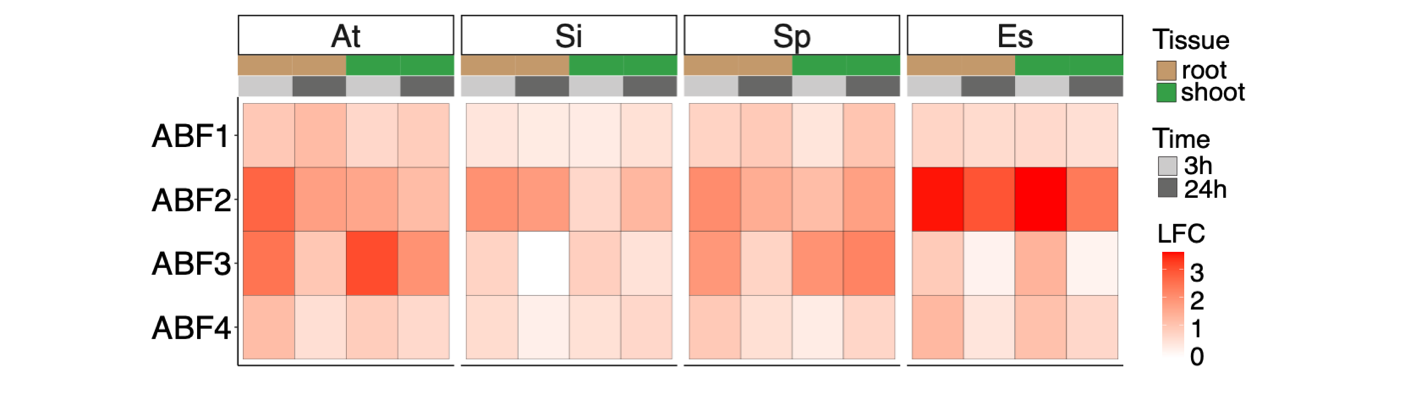

**fig. S5. Expression patterns of AREB/ABFs after ABA treatment.**

Heatmap shows the Log_2_ fold change values of AREB/ABFs 1, 2, 3, 4 upon ABA treatment at 2 separate tissue types (root and shoot) and timepoints (3 hours and 24 hours).

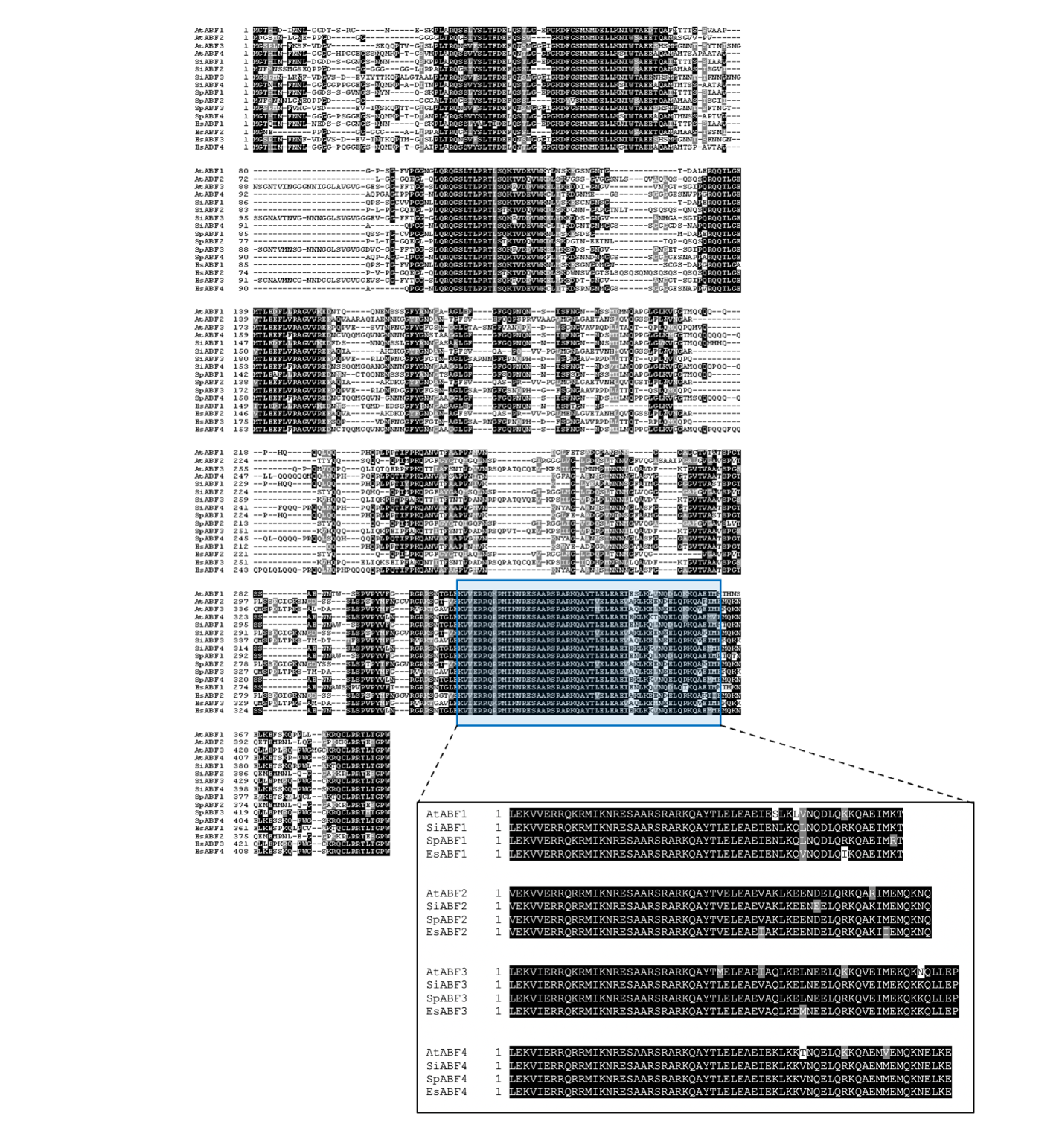

**fig. S6. AREB/ABFs of different species cluster together based on protein sequence and have conserved DNA-binding and protein-protein interaction domains.**

Alignment was generated using CLUSTALW with the amino acid sequences for each species. Black box highlights the conservation of the DNA binding domain and the protein-protein interaction domain for AREB/ABFs across species.

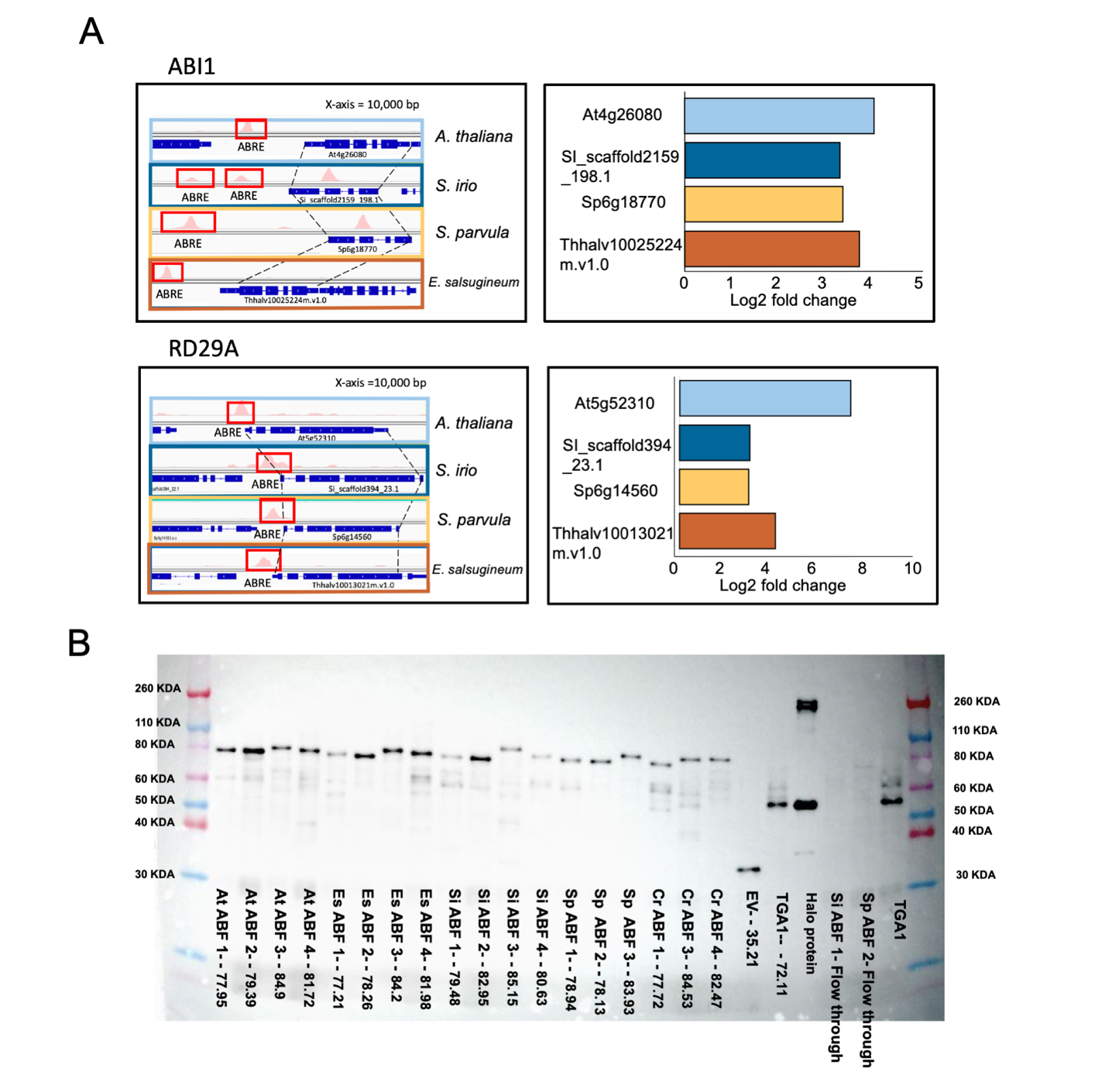

**fig. S7.** **Quality control checks to confirm the DAP-Seq assay works as expected.**

**(A)** DAP-Seq peaks associated with ABI1 and RD29A, which are marker genes for ABA response, are found across all species. Genome browser view shows binding sites represented as “peaks”. Dotted lines connect orthologous genes across species. ABREs are labeled in red and the data range for x-axis was standardized to 10,000bp. Corresponding differential gene expression data (ABA treatment vs. control from RNA-Seq for each gene ortholog across species is shown as log_2_ fold change. **(B)** Proteins synthesized by in vitro transcription/ translation (IVT) runs to expected size within typical yields as described in O’Malley *et al*. anti-HALO monoclonal antisera was used for detection of HALOtag used to generate the affinity matrix. GST-HALOTag (Halo protein) was used as positive control.

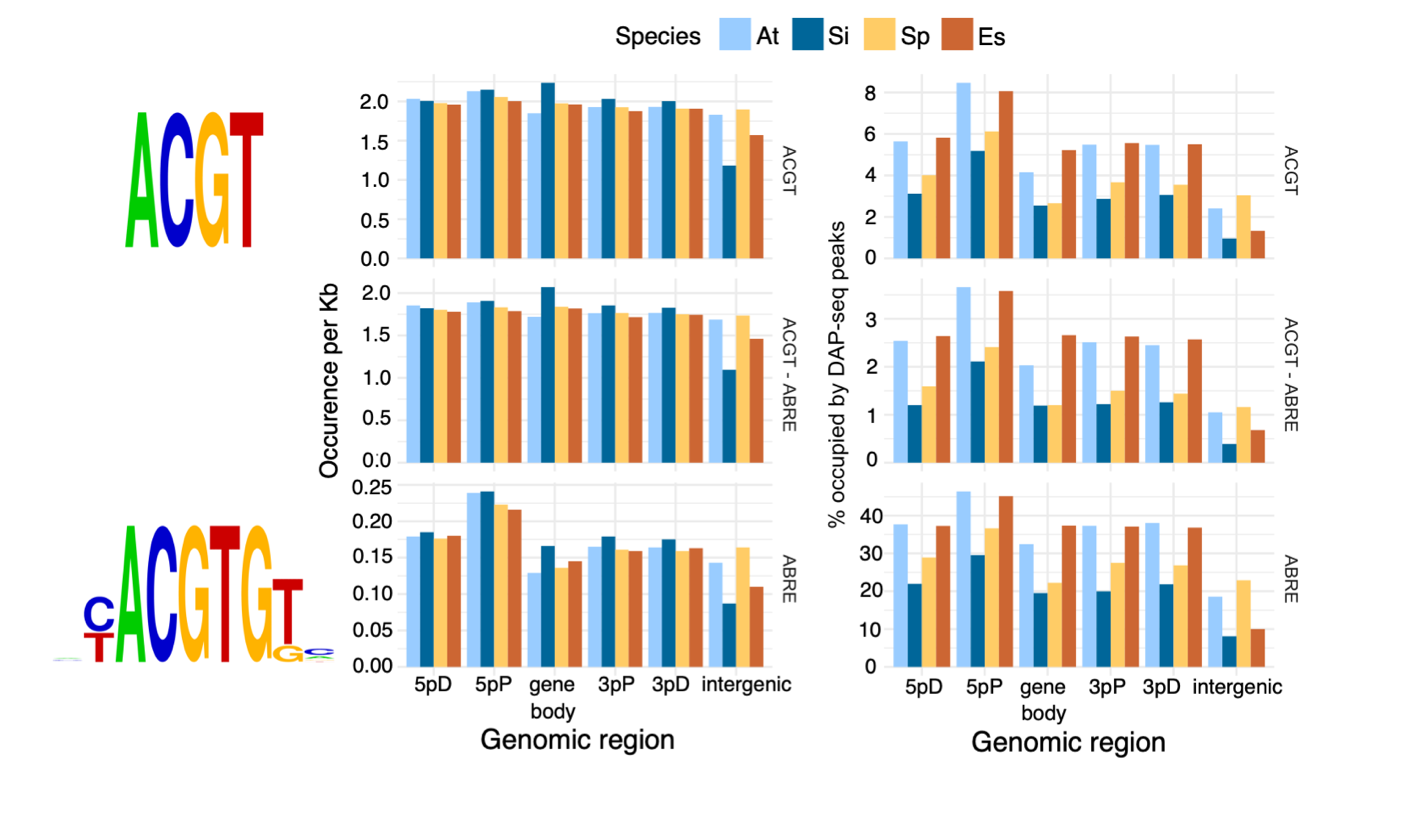

**fig. S8. Distribution of ACGT and ABRE motifs and proportions occupied by a DAP-seq peak.**

On the left panels, occurrences per kilobase (Kb) of the ACGT, ACGT sequences that are not a part of an ABRE (ACGT – ABRE), and ABRE motif, are plotted for different genomic regions adjacent to protein-coding gene models. Genomic regions are divided into 5’ distal (5pD), 5’ proximal (5pP), gene body, 3’ proximal (3pP), and 3’ distal (3pD), where proximal regions are within 1Kb of the coding sequence of a gene model and distal regions the next 1Kb blocks beyond the 1Kb regions. Intergenic regions are those not within 2Kb of any gene model. For each motif, percent proportions occupied by an AREB/ABF-binding DAP-seq peak are shown on the right panel. Motif logos indicate the definition of ACGT and ABRE motif. Genomic coordinates of all high-confidence DAP-seq peaks, together with their coincidence with ACGT or ABRE motifs and genomic regions adjacent to protein-coding gene models, are in data S6.

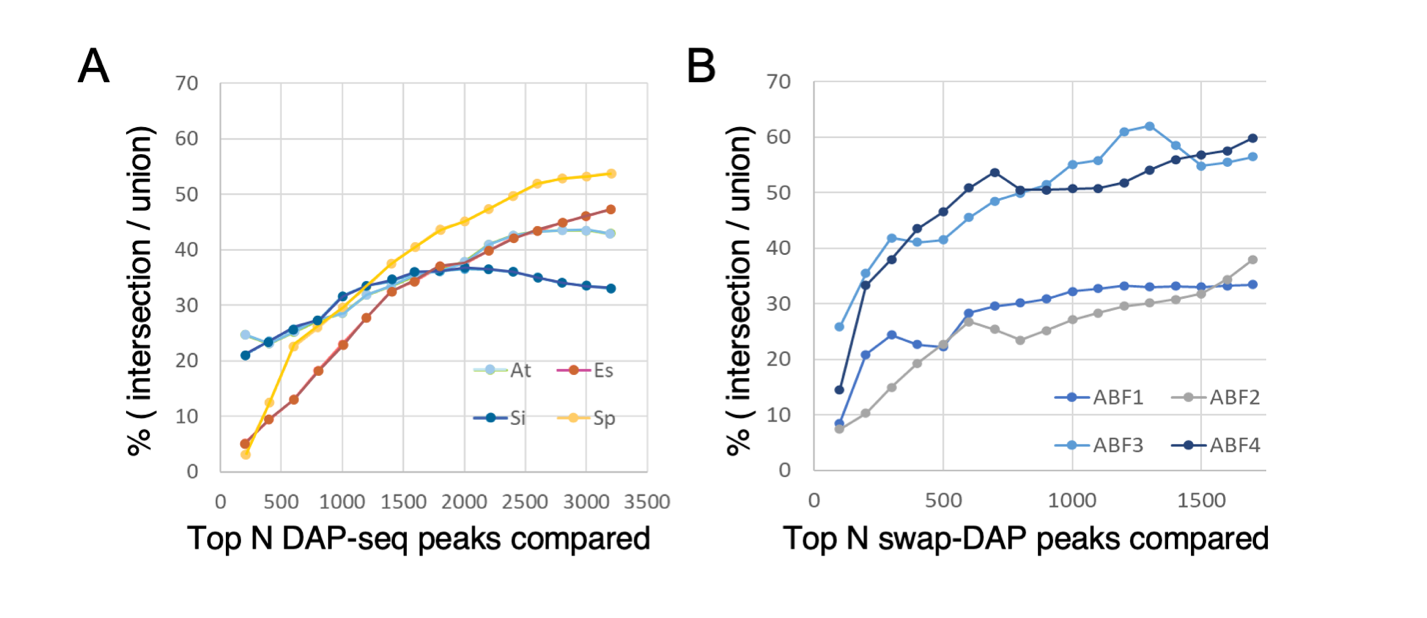

**fig. S9. Ranks and overlap of DAP-seq peaks.**

DAP-seq peaks were ranked based on significance and peak heights as detailed in Methods. We selected the top N ranked DAP-seq peaks from each experiment and compared their overlaps among AREB/ABF 1/2/3/4 within each species **(A)** or among AREB/ABF orthologs derived from *A. thaliana*/*S. irio*/*S. parvula*/*E. salsugineum* for each AREB/ABF in swap DAP-seq experiment. (**B**) Percentages of DAP-seq peak positions shared by all four AREB/ABFs (intersection) among all peak positions (union) were plotted over different N values

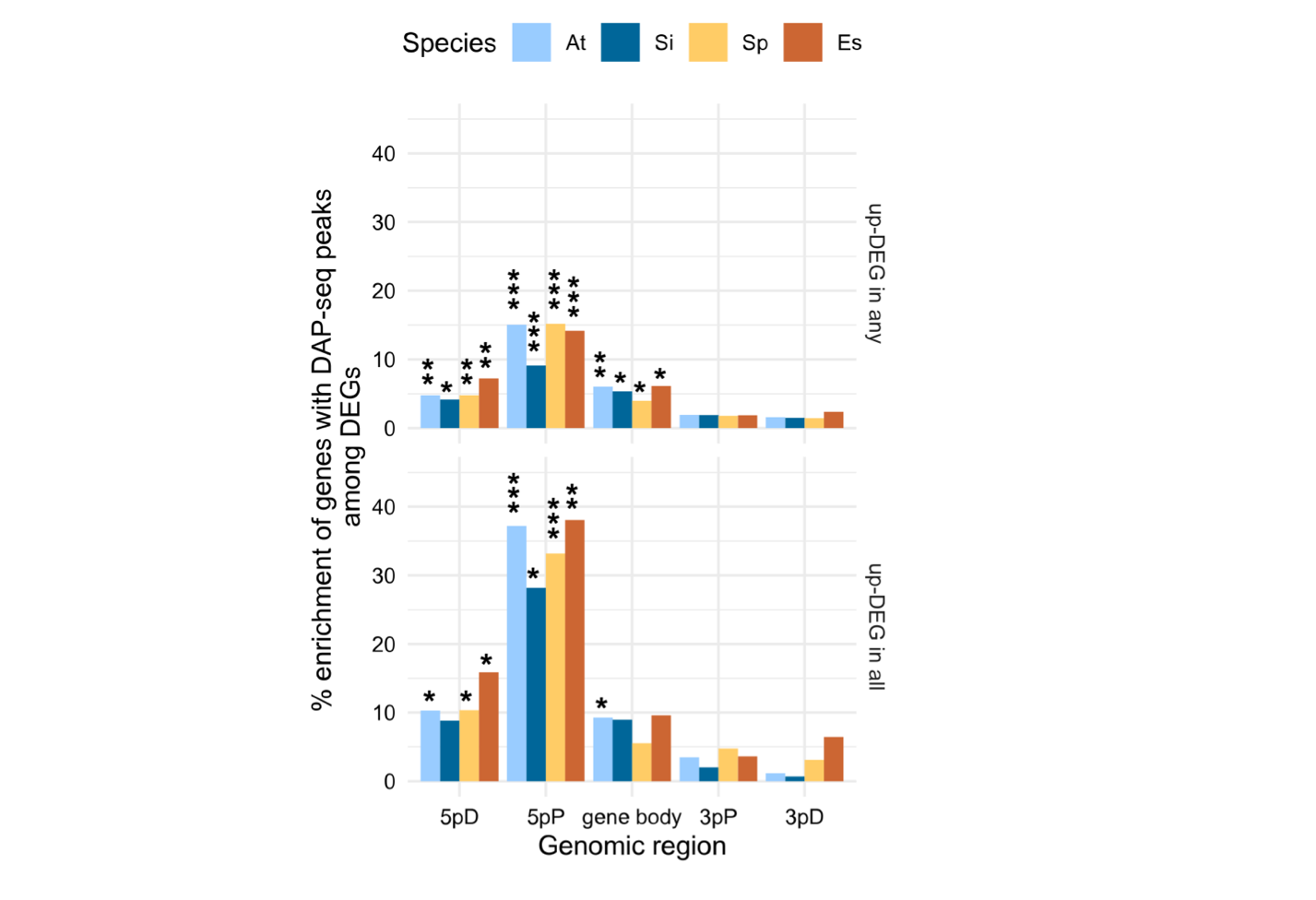

**fig. S10. Enrichment of genes associated with an AREB/ABF-binding DAP-seq peak among ABA-induced differentially expressed genes (DEGs).**

We tested whether genes including a DAP-seq peak in different genomic regions were enriched among ABA-induced DEGs compared to genes not regulated by ABA (background). Percent enrichment, compared to background, for genes significantly ABA-induced in any of the four samples (up-DEG in any sample, i.e. root or shoot tissues treated with ABA for 3hr or 24hr) and in all four samples (up-DEG in all samples) were plotted separately. Genomic regions are divided into 5’ distal (5pD), 5’ proximal (5pP), genomic coding sequences (gene body), 3’ proximal (3pP), and 3’ distal (3pD), where proximal regions are within 1Kb of the coding sequence of a gene model and distal regions the next 1Kb blocks beyond the proximal regions. Significance of enrichment (g-test with Benjamini-Hochberg correction) was marked for adjusted **P*< 10^-5^, ***P* <10^-20^, and ****P* <10^-50^. Significant enrichment of genes containing a DAP-seq peak in most genomic regions was not observed among ABA-repressed DEGs (table S2).

.

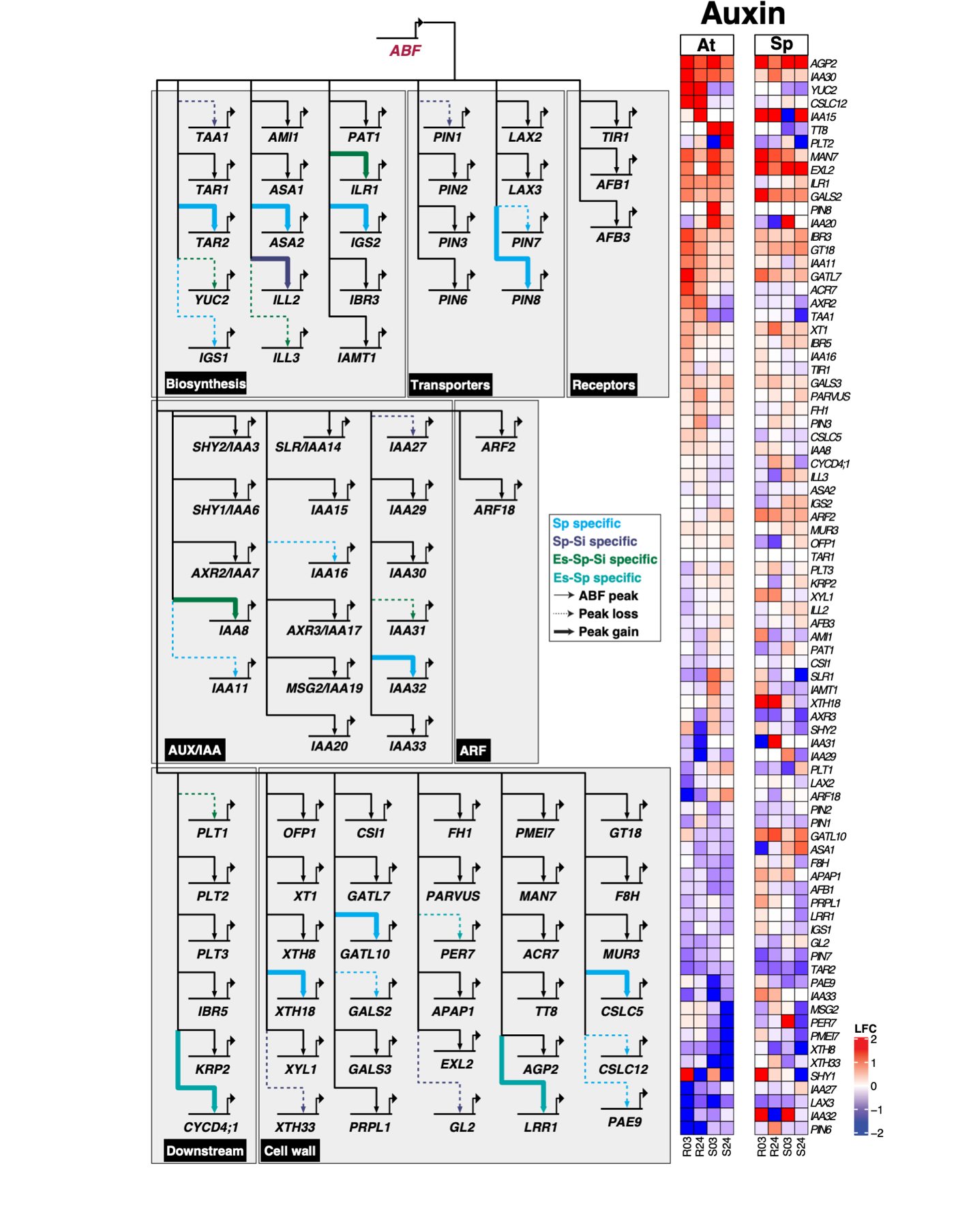

**fig. S11. Comparison between *A. thaliana* and *S. parvula* data sets reveal changes in AREB/ABF mediated auxin gene regulatory network.**

A total of 199 genes were curated for the analysis. 117 of the 199 genes related to primary auxin signaling in roots were present in our RNA-Seq data. 81 of these genes had AREB/ABF DAP peaks in either *A. thaliana* or *S. parvula* or in both suggesting AREB/ABF mediate gene regulation and shown in this GRN as nodes. Dotted arrow indicates loss of AREB/ABF binding, bold arrow indicates gain of AREB/ABF binding, and colors indicate differences that are lineage II specific, *S. parvula*-*S. irio*, *S. parvula*-*E. salsugineum*, or *S. parvula* specific differences compared to *A. thaliana*.

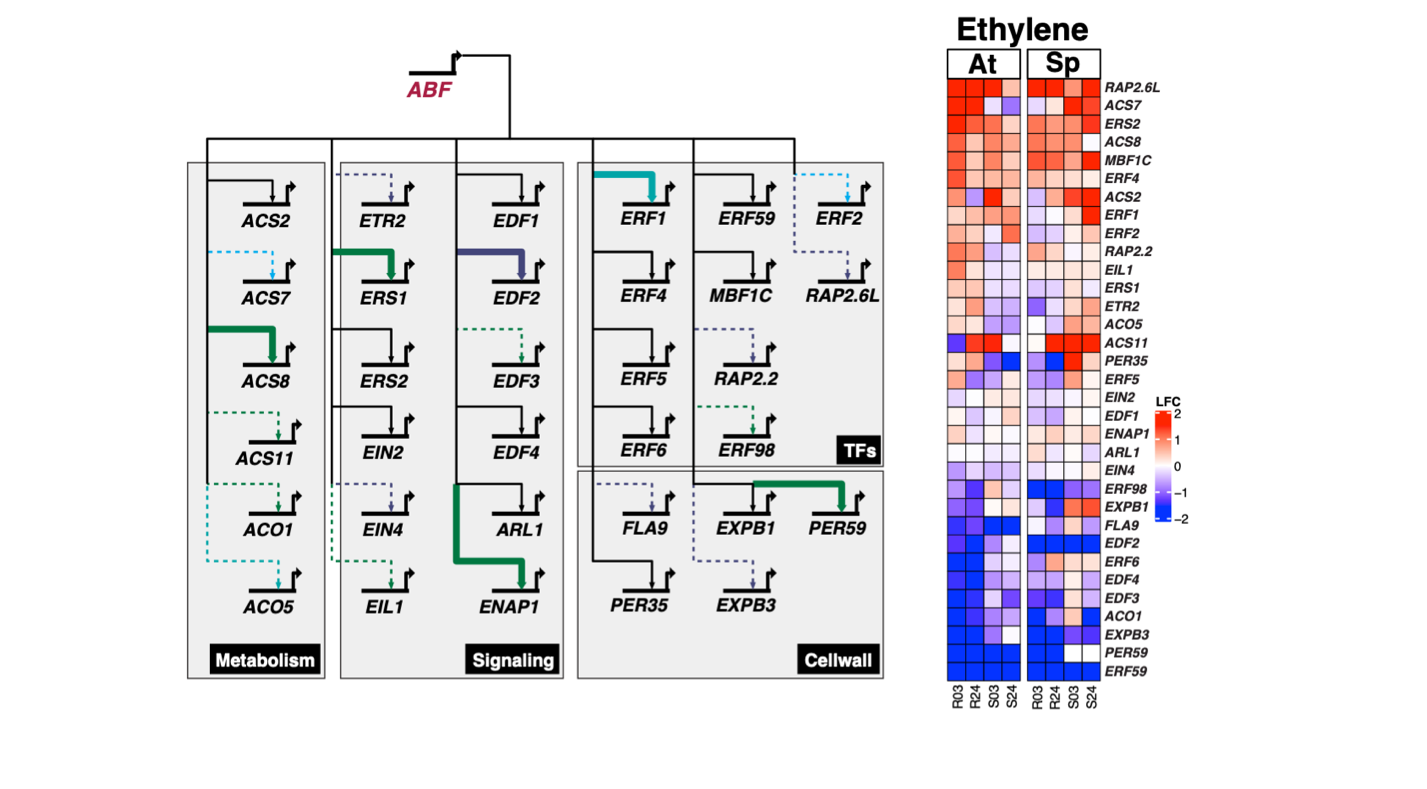

**fig. S12. Integration of AREB/ABF transcription factors into ethylene gene regulatory networks in *S. parvula*.**

Gene regulatory maps were drawn using BioTapestry, with a solid arrow line indicating correlation between AREB/ABF binding (DAP-Seq) and gene expression (RNA-Seq). An overlay of AREB/ABF-mediated GRN using ethylene regulatory network curated from previously published literature is shown. DAP-Seq and RNA-Seq data from different species were overlaid and differences are highlighted: dotted arrow indicates absence of AREB/ABF binding, bold arrow indicates presence of AREB/ABF binding, and colors indicate differences that are lineage II specific, *S. parvula*-*S. irio* specific, *S. parvula*-*E. salsugineum* specific, or *S. parvula* specific. Heatmap indicates differential gene expression patterns in all tissue types and time points for all nodes used to construct the GRN. Of the 72 genes identified, 50 were responsive to ABA, and 33 had AREB/ABF DAP peaks in either *A. thaliana*, *S. parvula*, or both which were used to construct the GRN. Among these however, only 2 were specific to *S. parvula* (ACS7 and ERF2) while the other 20 showed lineage II, *S. parvula*-*E. salsugineum*, or *S. parvula*-*S. irio* specific differences. Heatmap indicates differential gene expression patterns in all tissue types and time points for all nodes used to construct the GRN. LFC indicates log_2_ fold change.

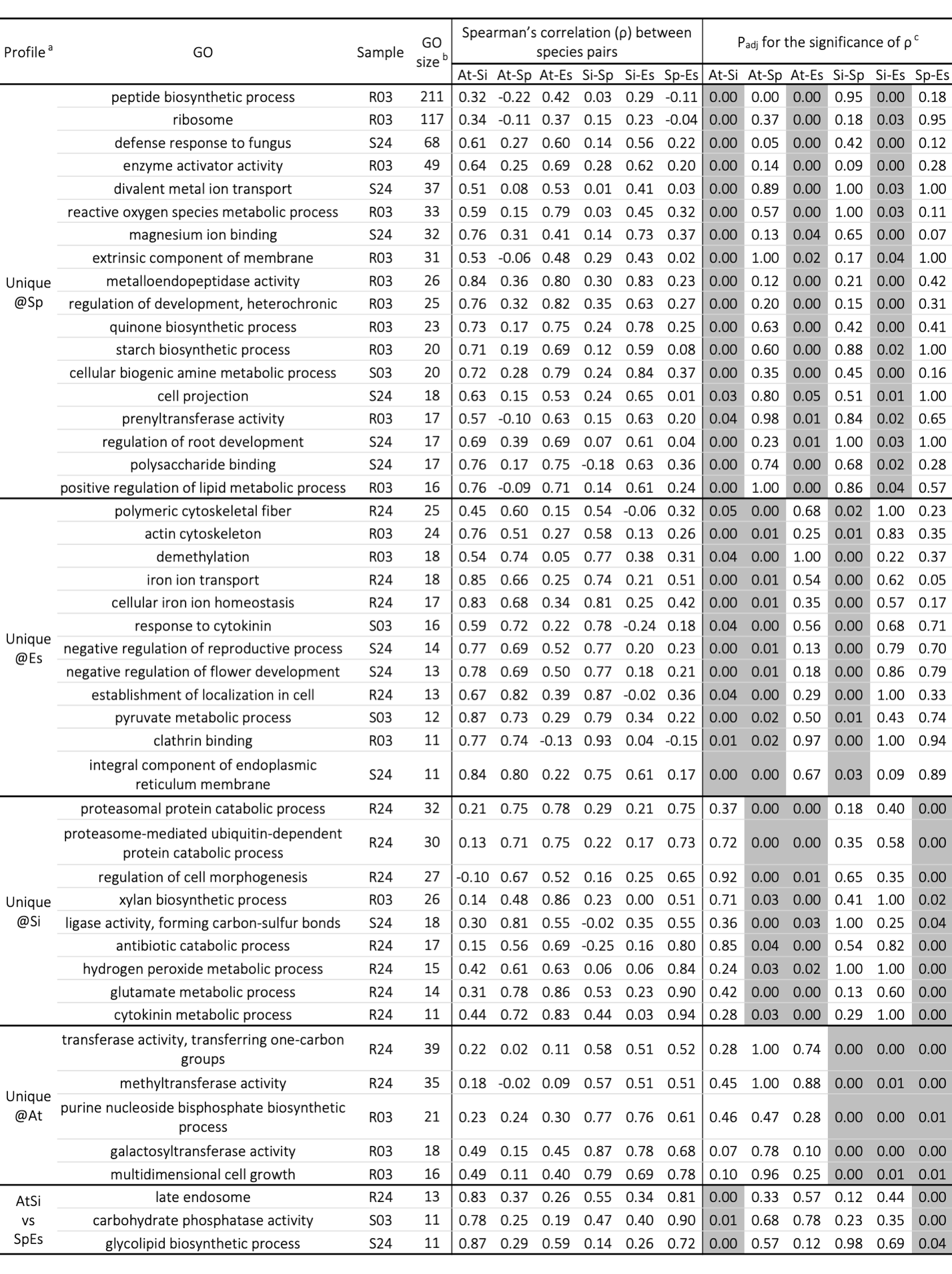
**table S1. Examples of GO terms showing lineage(s)-specific modification of ABA responsive expression.**

^a^ Phylogenetic profile “Unique@Sp,” “Unique@Es,” “Unique@Si,” and “Unique@At” are ABA responses unique in *S. parvula* (Sp), *E. salsugineum* (Es), *S. irio* (Si), and *A. thaliana* (At). Phylogenetic profile “AtSi vs. SpEs” indicates potential division between stress-sensitive (*A. thaliana* and *S. irio*) and extremophiles (*S. parvula* and *E. salsugineum*). ^b^ GO size (Gene Ontology size) contains the number of ortholog pairs annotated with the respective GO term. Median numbers among all six species pairs are shown, after filtering ortholog pairs as described in fig. S2. ^c^ Species pairs with significant positive Spearman’s correlations, based on adjusted *P*-value (P_adj_) < 0.05, are shaded in gray. Full results of the Phylogenetically informed profiling (Pip) analysis of ABA responses among the four Brassicaceae species are in data S4.

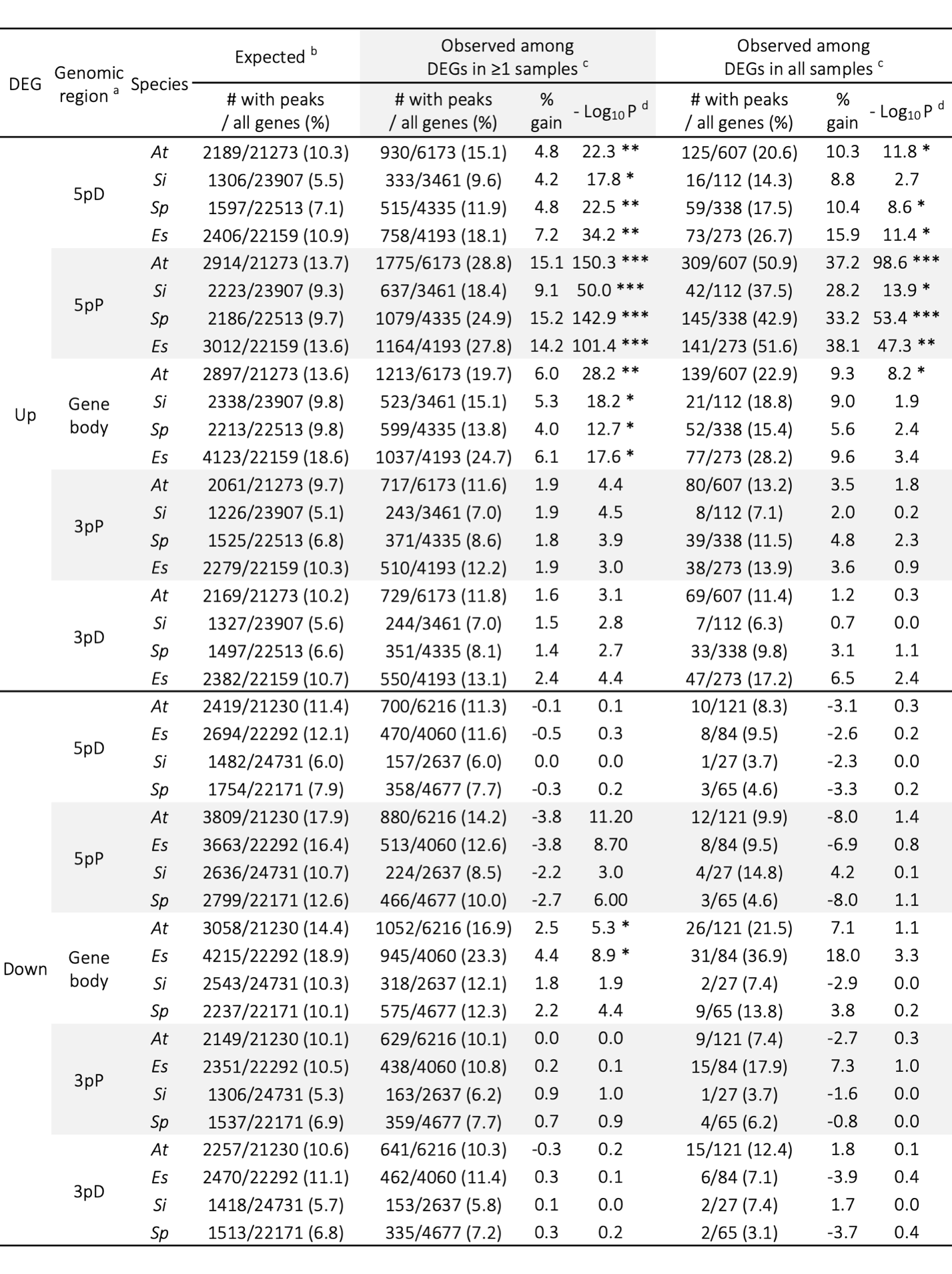

**table S2. Enrichment of genes associated with an AREB/ABF-binding DAP-seq peak among ABA-responsive differentially expressed genes (DEGs).**

^a^ Genomic region where an AREB/ABF DAP-seq peak was found is defined as: 5’ distal (5pD), 5’ proximal (5pP), genomic coding sequences (Gene body), 3’ proximal (3pP), and 3’ distal (3pD), where proximal regions are within 1Kb of the coding sequence of a gene model and distal regions the next 1Kb blocks beyond the proximal regions. ^b^ Proportion of genes with a DAP-seq peak among those not significantly differently expressed in any sample (i.e. in root and shoot tissues and 3hr and 24hr ABA treatment). ^c^ Values for genes differentially expressed in any of the four samples (≥1 samples) or all samples. ^d^ Based on g-test comparing the observed to the expected, with Benjamini-Hochberg correction for multiple testing. Adjusted *P*-value < 10^-5^ (*), <10^-20^ (**), and <10^-50^ (***) for positive enrichment.

**data S1. All 1-to-1 Orthologous groups with functional genomics data from RNA-seq and DAP-seq.**

ABA-responsive gene expression and number of DAP-seq peaks in adjacent genomic regions for 15,198 all unambiguous 1-to-1 ortholog groups.

**data S2. An analysis with OrthNet (including duplicates) with functional genomics data from RNA-seq and DAP-seq.**

An expanded version of data S1 where all genes, including lineage(s)-specific duplicates and those not in 1-to-1 ortholog groups, organized into OrthNet units based on both sequence similarity and co-linearity. See legend inside the Dataset for details.

**data S3. GO enrichment among ABA-induced and repressed DEGs.**

A matrix of GO terms enriched in ABA-induced and repressed DEGs for all species, tissues, and time points.

**data S4. Phylogenetically informed profiling (PiP) analysis of ABA-responsive gene expression for the four crucifer species.**

Results of PiP analysis identifying GO terms showing lineage(s)-specific modifications of ABA-responses. See legend inside the Dataset for details.

**data S5. JASPAR motifs enriched among promoters of ABA-induced and repressed DEGs.**

A matrix of known transcription factor-binding motifs, obtained from the JASPAR database, enriched in the promoters of ABA-induced and repressed DEGs for all species, tissues, and time points.

**data S6. All DAP-seq peak coordinates with annotations**

All DAP-seq peak coordinates with number of replicates, types of AREB/ABFs, and other information.

**data S7. Conserved ABA GRN.**

All genes curated to build the conserved ABA GRN shown in figure 4.

**data S8. Auxin and Ethylene GRN.**

All genes curated on the Auxin and ethylene pathway. 1:1 orthologs identified from RNA-seq and DAP-seq datasets were used to construct the actual GRN.
